# Data-driven biomarkers outperform theory-based biomarkers in predicting stroke motor outcomes

**DOI:** 10.1101/2023.06.19.545638

**Authors:** Emily R Olafson, Christoph Sperber, Keith W Jamison, Mark D Bowren, Aaron D Boes, Justin W Andrushko, Michael R Borich, Lara A Boyd, Jessica M Cassidy, Adriana B Conforto, Steven C Cramer, Adrienne N Dula, Fatemeh Geranmayeh, Brenton Hordacre, Neda Jahanshad, Steven A Kautz, Bethany Lo, Bradley J MacIntosh, Fabrizio Piras, Andrew D Robertson, Na Jin Seo, Surjo R Soekadar, Sophia I Thomopoulos, Daniela Vecchio, Timothy B Weng, Lars T Westlye, Carolee J Winstein, George F Wittenberg, Kristin A Wong, Paul M Thompson, Sook-Lei Liew, Amy F Kuceyeski

## Abstract

Chronic motor impairments are a leading cause of disability after stroke. Previous studies have predicted motor outcomes based on the degree of damage to predefined structures in the motor system, such as the corticospinal tract. However, such theory-based approaches may not take full advantage of the information contained in clinical imaging data. The present study uses data-driven approaches to predict chronic motor outcomes after stroke and compares the accuracy of these predictions to previously-identified theory-based biomarkers.

Using a cross-validation framework, regression models were trained using lesion masks and motor outcomes data from 789 stroke patients (293 female/496 male) from the ENIGMA Stroke Recovery Working Group (age 64.9±18.0 years; time since stroke 12.2±0.2 months; normalised motor score 0.7±0.5 (range [0,1]). The out-of-sample prediction accuracy of two theory-based biomarkers was assessed: lesion load of the corticospinal tract, and lesion load of multiple descending motor tracts. These theory-based prediction accuracies were compared to the prediction accuracy from three data-driven biomarkers: lesion load of lesion-behaviour maps, lesion load of structural networks associated with lesion-behaviour maps, and measures of regional structural disconnection.

In general, data-driven biomarkers had better prediction accuracy - as measured by higher explained variance in chronic motor outcomes - than theory-based biomarkers. Data-driven models of regional structural disconnection performed the best of all models tested (R^2^ = 0.210, p < 0.001), performing significantly better than predictions using the theory-based biomarkers of lesion load of the corticospinal tract (R^2^ = 0.132, p< 0.001) and of multiple descending motor tracts (R^2^ = 0.180, p < 0.001). They also performed slightly, but significantly, better than other data-driven biomarkers including lesion load of lesion-behaviour maps (R^2^ =0.200, p < 0.001) and lesion load of structural networks associated with lesion-behaviour maps (R^2^ =0.167, p < 0.001). Ensemble models - combining basic demographic variables like age, sex, and time since stroke - improved prediction accuracy for theory-based and data-driven biomarkers. Finally, combining both theory-based and data-driven biomarkers with demographic variables improved predictions, and the best ensemble model achieved R^2^ = 0.241, p < 0.001.

Overall, these results demonstrate that models that predict chronic motor outcomes using data-driven features, particularly when lesion data is represented in terms of structural disconnection, perform better than models that predict chronic motor outcomes using theory-based features from the motor system. However, combining both theory-based and data-driven models provides the best predictions.

## Introduction

Motor impairments are the most common type of deficit after stroke, and up to 50 percent of stroke survivors will have lasting hemiparesis.^1^ Providing accurate predictions of long-term motor outcomes is an ongoing goal of stroke research, as predictions based on acute clinical information can inform individualised rehabilitation strategies and can guide patient selection in clinical trials.^2,3^ Biomarkers derived from routinely-collected structural neuroimaging data that reflect lesion location with respect to critical white matter tracts have been related to motor outcomes,^4–8^ but there is no consensus on how to optimally model lesion damage to produce generalizable predictions of chronic motor deficits.

Historically, theory-based biomarkers, selected *a priori* based on their involvement in motor function, have been used to model motor outcomes after stroke. The most well-studied theory-based biomarker is the corticospinal tract (CST) lesion load, or the proportion of voxels in the ipsilesional corticospinal tract originating from primary motor cortex (M1) that intersect with the lesion.^9–13^ M1-CST lesion load has been related to motor deficits in the acute and chronic phase of stroke,^3,14^ but M1-CST damage in itself may not capture enough variance in lesion data to explain motor deficits in patients with a wide range of lesion topographies.^11,15,16^ Incorporating measures of damage to higher-order motor structures into linear models (e.g., lesion load of all tracts in the sensorimotor tract template atlas, SMATT-LL) helps to explain more variance in post-stroke motor outcomes compared to models based on measures of damage to M1-CST alone.^11,15–20^ Although lesion load to these tracts has been significantly associated with motor deficits within individual samples, the out-of-sample prediction performance of theory-based biomarkers has not been well-assessed.^14^

As an alternative to theory-based biomarkers, data-driven approaches assume that useful lesion-deficit associations can be discovered with sufficient data and proper representations of lesion damage.^21,22^ These approaches may be more suitable for predictive models than theory-based biomarkers: measures that are significantly related to motor outcomes in one sample may not predict outcomes in a new sample.^23^ Whether such data-driven approaches have value in stroke motor outcome prediction is unknown, and how to best represent lesion damage such that data-driven approaches can discover lesion-deficit associations that are predictive of motor outcomes is unclear. One approach is to discover voxels in which lesion damage is associated with motor deficits.^8,23^ The extent of overlap between a patient’s lesion and maps of lesion-deficit association - called lesion-behaviour maps (LBMs) - can be used as a biomarker in predictive models. Similarly, the extent of lesion overlap with structural lesion-network maps (sLNM), which reflect the white matter networks associated with peak LBM voxels, referred to previously as the sLNM lesion load,^8,23^ may be able to capture relationships between motor deficits and tract damage. One limitation of using voxelwise representations of damage to develop lesion-behaviour maps is that non-overlapping lesions that impact the same white matter tract are treated separately, which may reduce the power of a model to identify robust lesion-deficit associations.^24^ Transforming voxelwise lesions into structural disconnection measures may better represent the neural correlates of post-stroke deficits and improve statistical power to detect critical features.^24^ This type of representation is, in effect, a non-linear dimensionality reduction of voxelwise lesion data that can collapse damage to the same white matter tract by non-overlapping lesions into a single feature. To this end, the Network Modification tool^25^ can be used to calculate lesions’ Change in Connectivity (ChaCo) scores, reflecting the amount of structural disconnection to/from each grey matter region in the brain, by identifying white matter tracts that pass through the lesion using structural connectomes from healthy subjects.^25^

We hypothesised that data-driven biomarkers would outperform theory-based biomarkers in their ability to accurately predict chronic motor outcomes in held-out patients. Within data-driven biomarkers, we hypothesised that modelling lesion damage with whole-brain regional structural disconnection scores (ChaCo scores) would yield more accurate out-of-sample predictions of chronic motor scores than modelling lesion damage with lesion behaviour maps (LBM lesion load) and structural lesion network maps (sLNM lesion load), but that sLNM-LL would perform better than LBM-LL due to the inclusion of relevant structural networks.

Accurate, individualised outcome predictions inherently require that patient information is combined across several data sources. In addition to lesion damage, demographic factors such as age, sex, and time since stroke influence an individual’s chronic outcome. Models that employ combinations of imaging and demographic variables will likely be necessary for optimised predictions.^2^ Additionally, the strength of one imaging biomarker may be able to compensate for the weaknesses of others. Therefore, we hypothesised that predictive performance would be improved by incorporating demographic information and by combining predictions from several different biomarkers using ensemble models.

The variability between stroke subjects owing to the significant heterogeneity within the disease poses a challenge for clinical trials to identify effective therapies, presenting a need for biomarkers that are predictive of motor deficits and their ability to improve with treatment.^26,27^

## Materials and methods

### Sample demographics

A subset of cross-sectional data from the Enhancing Neuroimaging Genomics through Meta Analysis (ENIGMA) Stroke Recovery Working Group database (available as of 10 September 2021) from subjects with acute/subacute and chronic stroke was used in the study. Details of the ENIGMA Stroke Recovery procedures and methods are available in Liew et al.^28^ The data originated from 22 research studies carried out at different sites (Table 1). Informed consent was obtained from all subjects, and data were collected in compliance with each institution’s local ethical review boards and in accordance with the Declaration of Helsinki.

**Table 1:** Demographic information of the ENIGMA dataset, broken down by chronicity (acute/chronic) and by site within each chronicity group (some sites have both acute and chronic subjects that are listed separately). Total sample size (N), number of females (F) and males (M), and information about age (years), normalised motor scores, time since stroke at the time of assessment (months), and lesion volume (*cm*^3^). IQR, interquartile range

| Site ID | Total N.<br>N (F/M) | Median age<br>in years<br>(IQR) | Median motor<br>score (IQR) | Median time<br>since stroke<br>in mos. (IQR) | Median lesion vol.<br>in $cm^3$ (IQR) |
| --- | --- | --- | --- | --- | --- |
| **Acute +**<br>**subacute** |  |  |  |  |  |
| r005 | 1 (0/1) | 50.0 (0.0) | 0.29 (0.00) | 5.1 (0.0) | 1.69 (0.00) |
| r009 | 50 (13/37) | 70.0 (18.5) | 1.00 (0.05) | 0.2 (0.1) | 1.65 (6.96) |
| r025 | 9 (4/5) | 70.0 (19.0) | 1.00 (0.09) | 3.0 (1.0) | 0.52 (1.44) |
| r028 | 1 (0/1) | 63.0 (0.0) | 0.74 (0.00) | 5.5 (0.0) | 23.87 (0.00) |
| r031 | 36 (10/26) | 58.5 (13.2) | 0.52 (0.38) | 4.5 (1.7) | 10.83 (38.77) |
| r038 | 72 (30/42) | 66.5 (21.2) | 0.78 (0.66) | 2.9 (2.3) | 11.53 (41.64) |
| r040 | 57 (32/25) | 64.0 (22.0) | 0.40 (0.50) | 1.8 (1.5) | 14.72 (58.63) |
| r047 | 2 (1/1) | 71.0 (2.0) | 0.59 (0.26) | 4.4 (0.2) | 23.78 (19.19) |
| r049 | 21 (12/9) | 65.0 (16.0) | 0.95 (0.00) | 0.0 (0.0) | 1.30 (2.18) |
| r050 | 14 (7/7) | 68.0 (16.8) | 0.92 (0.10) | 0.0 (0.0) | 0.33 (0.40) |
| r053 | 52 (20/32) | 63.5 (20.5) | 0.92 (0.17) | 3.0 (3.0) | 13.50 (28.27) |
| r054 | 12 (5/7) | 65.5 (11.8) | 0.67 (0.83) | 0.4 (0.2) | 4.06 (14.15) |
| **Chronic** |  |  |  |  |  |
| r001 | 39 (10/29) | 61.0 (17.0) | 0.65 (0.23) | 23.5 (40.0) | 6.27 (18.06) |
| r002 | 12 (6/6) | 69.5 (11.5) | 0.50 (0.41) | 73.2 (51.9) | 28.24 (31.71) |
| r003 | 15 (6/9) | 61.0 (16.5) | 0.24 (0.20) | 48.8 (67.6) | 20.28 (76.88) |
| r004 | 19 (7/12) | 44.0 (14.5) | 0.17 (0.16) | 50.4 (81.9) | 36.85 (44.29) |
| r005 | 27 (12/15) | 66.0 (16.5) | 0.79 (0.45) | 31.4 (27.8) | 1.61 (40.27) |
| r009 | 60 (17/43) | 71.0 (7.2) | 0.96 (0.12) | 27.4 (9.3) | 1.43 (4.65) |
| r025 | 16 (3/13) | 64.5 (13.2) | 0.98 (0.58) | 14.2 (10.2) | 5.92 (14.26) |
| r027 | 28 (8/20) | 57.0 (10.2) | 0.30 (0.16) | 19.3 (24.7) | 12.30 (62.13) |
| r028 | 21 (6/15) | 63.0 (9.0) | 0.82 (0.24) | 26.5 (37.5) | 5.25 (41.28) |
| r031 | 1 (0/1) | 52.0 (0.0) | 0.68 (0.00) | 6.1 (0.0) | 1.54 (0.00) |
| r034 | 15 (6/9) | 58.4 (11.1) | 0.82 (0.20) | 61.3 (68.3) | 6.68 (34.99) |
| r035 | 15 (6/9) | 64.0 (18.0) | 0.64 (0.52) | 33.5 (22.9) | 3.89 (31.56) |
| r038 | 18 (7/11) | 67.0 (10.0) | 1.00 (0.12) | 15.1 (10.1) | 1.98 (1.63) |
| r040 | 14 (7/7) | 63.5 (9.8) | 0.68 (0.47) | 14.1 (17.5) | 8.65 (82.65) |
| r042 | 22 (11/11) | 48.5 (15.5) | 0.64 (0.19) | 29.6 (36.4) | 14.16 (49.53) |
| r044 | 4 (0/4) | 68.0 (9.2) | 0.52 (0.25) | 43.7 (52.9) | 23.65 (67.00) |
| r045 | 4 (1/3) | 62.0 (5.2) | 0.49 (0.24) | 96.1 (59.0) | 7.97 (6.66) |
| r046 | 11 (3/8) | 62.0 (10.5) | 0.50 (0.29) | 86.3 (83.4) | 4.62 (19.82) |
| r047 | 44 (14/30) | 65.5 (12.0) | 0.65 (0.44) | 38.1 (53.7) | 12.72 (41.33) |
| r048 | 43 (16/27) | 68.0 (12.5) | 0.79 (0.44) | 46.2 (49.8) | 7.93 (43.45) |
| r052 | 32 (12/20) | 63.0 (13.5) | 0.41 (0.09) | 39.1 (42.2) | 6.98 (51.55) |
| **All** |  |  |  |  |  |
|  | 789 (293/496) | 64.0 (18.0) | 0.7 (0.5) | 12.2 (0.2) | 6.45 (32.48) |

ENIGMA Stroke Recovery subjects with the following data were included: (1) high-resolution (1-mm isotropic) T1-weighted brain MRI (T1w) acquired with a 3T MRI scanner; (2) information about time since stroke at time of imaging, as well as (3) age, (4), sex, and (5) a measure of sensorimotor function from one of the following assessments: i) Fugl-Meyer Assessment of Upper Extremities (FMA-UE), a performance-based measure of paretic upper extremity impairment,^29^ ii) the Barthel index, which measures the extent to which a person can function independently and has mobility in their activities of daily living,^30^ or iii) the National Institutes of Health Stroke Score (NIHSS), a broad measure of stroke severity that includes assessment of non-motor and motor functions.^31^ For most subjects with chronic stroke, motor deficits are normalised FMA-UE scores, whereas normalised FMA-UE scores were available for fewer of the subjects with acute/subacute stroke (Supplementary Table 1); for simplicity, we refer to all outcomes as ‘motor’ scores, though the main models were replicated with a single motor assessment (Fugl Meyer UE scores) as well. Motor scores were normalised to the range [0, 1] by dividing the raw score by the maximum possible score for that assessment. Behavioural data were collected within approximately 72 hours of the MRI. Subjects were considered in the chronic phase of stroke if their time since stroke at the time of assessment was greater than or equal to 180 days, and were considered to be in an acute/subacute phase if they were assessed within 180 days after the stroke. See Supplementary Fig. 1 for lesion distribution in MNI space.

### General overview

We built several models to predict chronic motor scores from imaging data and minimal demographic information. Each model used lesion-derived information as input to predict normalised motor scores in subjects with chronic stroke. We compared several different biomarkers that reflect different aspects of lesion damage. These biomarkers include:

### Theory-based biomarkers

▪ M1-CST LL (1 feature)
▪ SMATT-LL (6 features for ipsilesional tracts, 12 features for bilateral tracts)

### Data-driven biomarkers

▪ LBM-LL (1 feature)
▪ sLNM-LL (5 features; 5 components derived from principal components analysis of structural connectivity seeded from lesion-behaviour map)
▪ ChaCo scores (86 or 268 features depending on the atlas, fewer when using feature selection)

Nested cross-validation was performed during model training and performance was assessed on unseen test data. For ChaCo score models, we assessed whether feature selection improved performance. Additionally, we evaluated whether including acute subjects in the training set, but not in the test set, improved prediction of chronic deficits, with the idea that the larger variance in motor scores in the acute data would be useful in predicting motor deficits in the chronic phase. We also evaluated whether adding basic demographic information (age, sex, time since stroke) via ensemble models improved performance. Finally, we assessed whether using ensemble models to combine predictions from multiple different lesion damage metrics improved performance.

### Machine learning framework

Regression models were trained and evaluated using repeated 5-fold nested cross-validation (Figure 1). Model implementation differed for each lesion biomarker based on the dimensionality of the data; see below for implementation details for each biomarker (section: Description of models and their inputs). For biomarkers with more than one feature, ridge regression models were fit with a single hyperparameter indicating the degree of regularisation. In the outer loop, the data were split into 5 training and test partitions. First, models were trained using all data (including acute, subacute, and chronic timepoints), and tested only on chronic subjects, yielding 696 subjects in the training set and 92 subjects in the test set. Using only chronic data to train the models, there were approximately 370 subjects in the training set and 92 subjects in the test set. Out-of-sample performance was calculated as the average performance across 5 outer test folds. We obtained a distribution of out-of-sample performance by splitting the data into 5 train/test folds 100 times, shuffling the indices of the splits each time.

**Figure 1.**
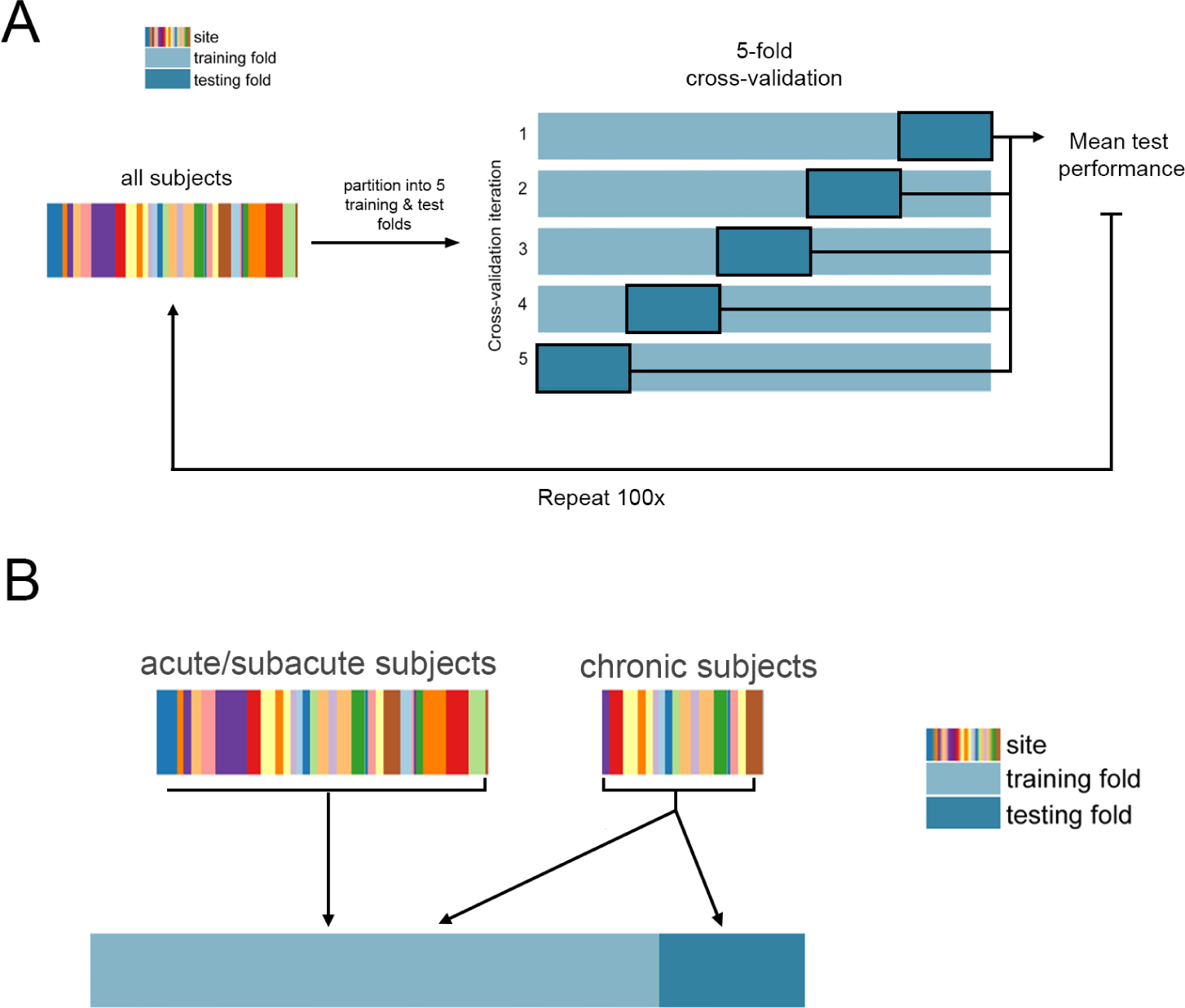
Cross-validation framework. **A**. Overview of 5-fold cross-validation. Subject data is partitioned into five non-overlapping training and test folds, such that no training subjects are in the test set, and no subject is in the test fold more than once. **B.** Use of acute/subacute subjects in training folds but not test folds. When using all training data, chronic subjects were included in the test folds and training folds, whereas acute/subacute stroke subjects were only included in training folds.

### Replication with Fugl-Meyer Upper Extremity Assessments only

Although the majority of the chronic stroke subjects had FMA-UE scores as their outcome measure, the acute/subacute stroke subjects had more varied assessments, including measures that describe more general functional outcomes like the Barthel index and NIHSS. Although these measures can be correlated with motor endpoints, they are influenced by many factors beyond motor status. We therefore replicated the main models using a subset of the data with FMA-UE scores (N subset = 392) to compare data-driven versus theory-based biomarkers on their ability to predict a single motor measure.

### Model performance

Model performance was assessed by comparing true normalised motor scores with predicted scores. Performance was calculated with both Pearson’s correlation coefficient and explained variance, or R^2^, which captures the percent of variation in motor scores explained by variation in the model predictors:

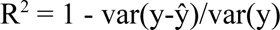

Where y is a vector of true motor outcomes, and ŷ is a vector of predicted motor outcomes. These two performance metrics were used to compare results with prior literature, but differences between models were assessed using R^2^, as it is a more robust metric to assess model quality.

Differences in performance between models were assessed using two-sided Wilcoxon signed-rank tests and p-values were corrected for multiple comparisons using Bonferroni correction (p < 0.05).

For each predictive model, we generated a null distribution for assessing model significance by permuting the predicted variable (motor score) 100 times. Then, as for each normal model, we ran 100 5-fold train/test cross validation splits for each permutation. This yielded a distribution of 100 out-of-sample mean performance measures for each permutation. The median across these 100 measures was then calculated for each permutation. In total, 100 null median performance measures were calculated. The p-value for the model’s significance is the proportion of null models that had median prediction accuracies greater than or equal to the median performance of the true model.

### Ensemble models

The idea of ensemble learning is to build a single prediction model by combining the strengths of a collection of simpler base models; we used ensemble models that average predictions from different biomarkers.^32^ We tested whether models including demographic information (age, sex, and days post stroke), ensembled with lesion models, performed better than models with lesion data or demographic data alone (i.e., we hypothesised that the variance explained from lesion data and demographic data is not redundant). We also assessed whether models including both lesion load and ChaCo scores would perform better than models with lesion load or ChaCo scores alone. Ensemble models were generated by training ChaCo-models and lesion load models separately, on the same subjects and with the same training/test/validation splits, and averaging the final predicted scores for each test subject. Standard linear regression was used to model the relationship between demographic information and motor impairment.

### Analysing feature weights

Feature weights in high-dimensional models can be unstable and therefore only provide limited interpretability.^33^ To assess the robustness of feature weights (i.e., beta coefficients), the Pearson correlation in regional feature weights across all training folds was calculated. For this specific feature stability analysis (but not for model evaluation), acute/subacute subjects were split for each training fold such that different folds did not contain the same set of acute/subacute subjects (all acute/subacute subjects were included in all training folds in the model evaluation phase to maximise the amount of data available for training). For the 86-region ChaCo models, the average of the 86-region *β* vector, representing the model weights for each region, across 5 folds was calculated for each of the 100 train/test splits. The median across these 100 splits was visualised on a glass brain. For the 268-region ChaCo models with feature selection, the most consistently-selected regions (selected in at least 475/500 folds or 99% of outer folds) were identified. The average of the 268-region *β* vector across the 5 folds was calculated for each of 100 train/test splits. The median average *β* weight for consistently-selected regions across these 100 splits was visualised on a glass brain.

## Description of models and their inputs

### Primary motor cortex CST lesion load (M1-CST-LL) models

The lesion load of the corticospinal tract originating from the primary motor cortex (M1-CST-LL) was calculated (Figure 2A). Here, as in previous work, M1-CST-LL was calculated as the proportion of lesioned voxels intersecting with a binarized ipsilesional M1-CST template.^9^ Specifically, lesion load was calculated in 1-mm MNIv6 space as:

**Figure 2.**
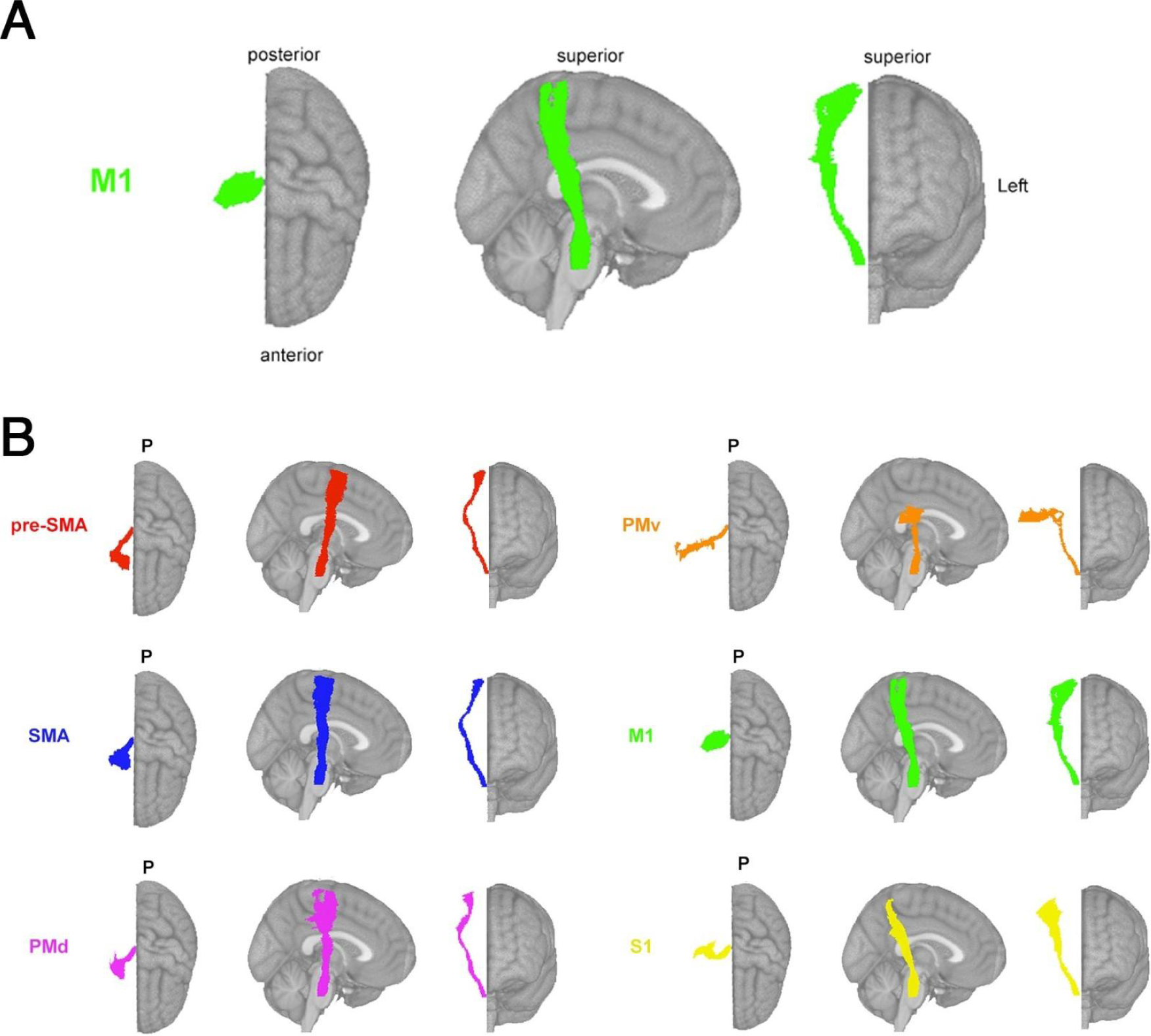
Theory-based biomarkers. **A**. The M1-CST, here displaying only right hemisphere tracts relative to an MNI template. **B.** Tracts from the sensorimotor tract template atlas (SMATT), displaying only right hemisphere tracts relative to an MNI template, including pre-supplementary motor area (pre-SMA), supplementary motor area (SMA), dorsal premotor cortex (PMd), ventral premotor cortex (PMv), primary motor cortex (M1), and primary sensory cortex (S1). Pre-SMA is the most anterior tract, S1 is the most posterior tract.

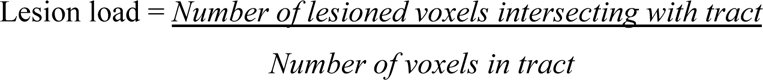

Left and right hemisphere M1-CST segmentations in MNI space were obtained from the high-resolution sensorimotor area tract template (SMATT).^34^ Few subjects had non-zero M1-CST-LL values (Supplementary Fig. 2A). Linear regression was used to model the relationship between ipsilesional M1-CST-LL and chronic motor scores. The weights from the best-performing model in the inner loop were used to predict motor scores for new subjects in the test folds.

### Sensorimotor tract lesion load (SMATT-LL) models

Sensorimotor tract segmentations were obtained from the sensorimotor area tract template (SMATT),^34^ which contains 12 tracts derived from probabilistic tractography seeded in the left and right primary motor cortex (M1), dorsal and ventral premotor cortex (PMd and PMv, respectively), supplementary motor area (SMA), pre-supplementary motor area (pre-SMA), and primary somatosensory cortex (S1) performed in healthy controls (Figure 2B). Lesion load was calculated as above for all 12 bilateral tracts (L/R SMATT-LL) and for 6 ipsilesional tracts (ipsilesional SMATT-LL). L/R SMATT-LL was calculated in order to assess whether preserving hemispheric information improved predictions. For subjects with brainstem, cerebellar, and/or bilateral cerebral strokes, ipsilesional lesion load was calculated as the average lesion load of the left and right hemisphere tracts (Supplementary Fig. 2B,C).

Ridge regression models were used to predict chronic motor deficits from ipsilesional SMATT-LL (6 features) and from L/R-SMATT-LL (12 features). Ridge regression was used to account for multicollinearity of lesion load values between tracts (Supplementary Fig. 3, 4). Lesion load values were normalised (after train/test split) by subtracting the mean across subjects and dividing by the l2-norm prior to model fitting. In the inner loop, the degree of model regularisation (*λ*) was determined via grid-search over 30 values ranging from 10*^−^*^2^ to 102. The training data was fit with the selected *λ* and this model was used to predict motor scores for held-out subjects in the test folds.

### Lesion-behaviour map lesion load (LBM-LL) models

A lesion behaviour map (Figure 3A) was obtained as described by Bowren et al. (2022). Specifically, Bowren et al. used sparse canonical correlation analysis to produce maps of voxels in which damage was associated with Fugl-Meyer scores.^35^ Lesion load to this lesion-behaviour map (LBM-LL) was calculated as the sum of voxels in the LBM that intersect with the lesion. Standard linear regression models were used to predict chronic motor deficits from LBM-LL.

**Figure 3.**
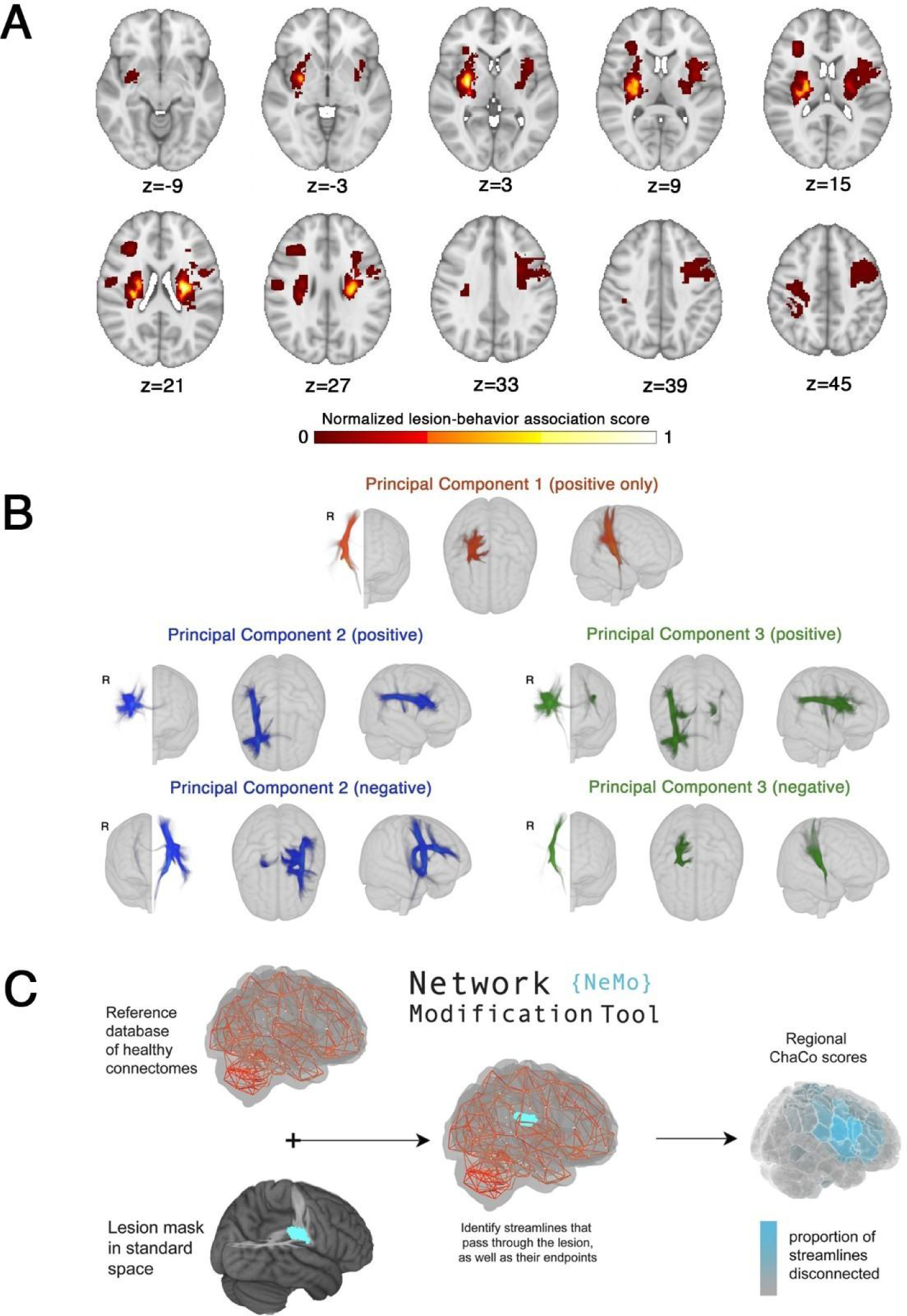
Data-driven biomarkers. **A**. Lesion-behaviour map (LBM) representing the association between voxelwise damage and Fugl-Meyer scores, derived from multivariate lesion-behaviour mapping with Fugl-Meyer scores. **B.** Structural lesion network maps (sLNMs), derived from seed-based tractography run on peak regions identified from LBM (A) and then performing principal components analysis to identify 3 components, split into positive and negative weights. **C.** Change in Connectivity (ChaCo) scores derived from the Network Modification (NeMo) tool. Binary lesion masks in MNI space representing the presence of a stroke lesion (turquoise) in a given voxel are provided by the user. Each lesion mask is embedded into 420 unrelated healthy structural connectomes (separately for each healthy subject) and the regional ChaCo scores are calculated and averaged across healthy subjects (parcellation shown here is the Shen 268-region atlas)

### Structural lesion-network mapping lesion load (sLNM-LL) models

Structural lesion network maps (Figure 3B) were obtained from Bowren et al. (2022). Specifically, peak white matter (WM) voxels from lesion-behaviour maps (described above) were identified. Then, tractography was seeded from these peak WM voxels to identify associated structural networks, called structural lesion network maps (sLNMs). Principal components analysis of sLNMs was performed, which produced 3 principal components that correspond to 5 sLNM maps (PC1, and positive/negative weights of PC2 and PC3). Lesion load on each sLNM map was calculated for each subject as the sum of the voxel intensities from the principal component map that intersected the lesion mask (Supplementary Fig. 5). Ridge regression models were used to predict chronic motor deficits from sLNM lesion loads (5 features). As above, lesion load values were normalised (after train/test split) by subtracting the mean across subjects and dividing by the l2-norm prior to model fitting. Hyperparameter optimization was performed as described above for SMATT models.

### Regional change in connectivity (ChaCo) models

Lesion masks in 1-mm^3^ MNI v6 space were processed with the Network Modification Tool (NeMo Tool) v2 pipeline,^25^ available at https://kuceyeski-wcm-web.s3.us-east-1.amazonaws.com/upload.html; see https://github.com/kjamison/nemo for documentation. Given a lesion mask, the NeMo tool produces outputs that reflect the impact of the lesion on white matter tracts using healthy structural connectomes as a reference. The NeMo tool embeds a lesion mask into healthy structural connectomes, identifies all white matter streamlines that intersect with the lesion, and determines the brain regions at the endpoints of those streamlines (Figure 3C). Regional change in connectivity (ChaCo) scores, or the ratio of the number of disrupted streamlines divided by the total number of streamlines terminating in each region, were calculated for all grey matter regions (see Supplementary Fig. 6A,B for distribution of mean and standard deviation of ChaCo scores). The NeMo tool uses structural connectivity from 420 unrelated subjects from the Human Connectome Project (HCP) Young Adult database. Regional ChaCo scores from two different atlases were compared: the 86-region Desikan-Killiany Atlas (68 cortical regions + 18 subcortical regions, excluding brainstem) from FreeSurfer (“fs86” for short), which contains coarse anatomically parcellated regions,^36,37^ and the 268-region Shen atlas (“shen268” for short), which contains more fine-grained functionally parcellated cortical and subcortical regions.^38^

First, the performance of ridge regression models was assessed, as described above, with regional ChaCo scores as inputs (86 features for the fs86 atlas, 268 features for the shen268 atlas). Then, a filter-based feature selection step was added to the ridge regression models to obtain a subset of features that were the most useful for prediction.^39^ Features were ranked by their association with the outcome variable (p-value from univariate regression) and only the *κ* most associated variables were included in the model. In the inner hyperparameter selection loop, both the amount of regularisation on regression coefficients (*λ*) and the number of features to retain in the model (*κ*) were selected via grid search. The *λ* value was chosen by searching over 30 values ranging 10*^−^*^2^ to 10^2^, and the *κ* value was chosen by searching 30 values ranging from 5 to the maximum number of features possible (for fs86: 86, for shen268: 268). This feature selection approach was implemented to identify sparse sets of correlated variables, as opposed to more basic embedded feature selection techniques such as LASSO^40^ that randomly suppresses collinear features.

## Code availability

The scikit-learn package was used to implement machine learning models (http://scikit-learn.org). All analysis scripts that generated the results of the present study are available as open source (https://github.com/emilyolafson/lesion_predictions), and the LBM and sLNM maps are also available on the repository. The script can be easily modified to predict any outcome score from ChaCo scores/lesion load data.

## Data availability

To protect the privacy of research participants, individual subject data used in this study are not available in a public repository. Participating research cohorts vary in public data-sharing restrictions as determined by the following: (1) ethical review board and consent documents; (2) national and transnational sharing laws; and (3) institutional processes that may require signed data transfer agreements for limited, predefined data use. However, data sharing is possible for new and participating ENIGMA Stroke Recovery Working Group members who agree to the consortium’s ethical standards for data use and upon the submission of a secondary analysis plan for group review. Upon the approval of the proposed analysis plan, access to relevant data is provided contingent on local principal investigator approval, data availability, and compliance with supervening regulatory boards. Deidentified summary data used for this study can be made available upon reasonable request to the corresponding author.

## Results

### Relative performance of models

The out-of-sample performances of the models using all training data can be found in Figure 4A,B. All models performed significantly better than chance (p < 0.001). With the exception of sLNM-LL models, all data-driven models (i.e., LBM-LL and ChaCo models) outperformed all theory-based models when using all training data (Figure 5A). When using only chronic data for training, only LBM-LL models outperformed all theory-based models (Figure 5B, Supplementary Fig. 7A,B).

**Figure 4.**
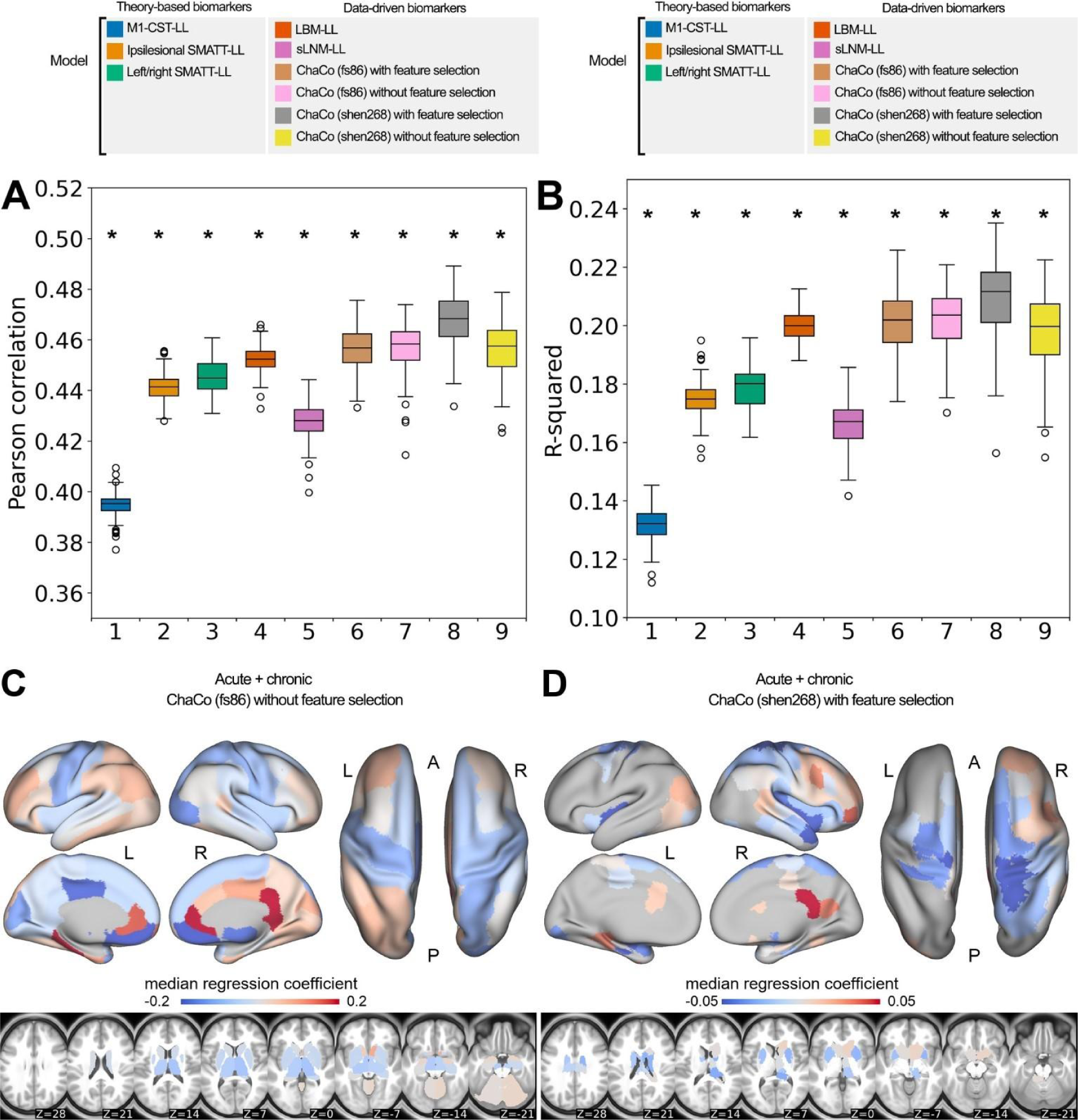
Summary of model performance metrics across all models tested and feature weights (regression coefficients *β*) for the two best-performing models. **A.** and **B.** Distribution of model performance (mean Pearson correlation/*R*^2^ across 5 outer folds for 100 permutations of the data). Asterisks (*) indicate that model performance is significantly above chance (\**, p <* 0.001), as assessed via permutation testing. The boxes extend from the lower to upper quartile values of the data, with a line at the median. Whiskers represent the range of the data from [Q1-1.5*IQR, Q3+1.5*IQR]. **C.** and **D.** Mean feature weights for the top two best-performing models (ChaCo (fs86) without feature selection, ChaCo (shen268) with feature selection, respectively). For the fs86-ChaCo model (left), we display the mean regression coefficients *β* across 100 permutations. For ChaCo (shen268) (right), we display the median regression coefficients of regions that were selected in at least 95% of outer folds (i.e., for regions that were included in the model in at least 475/500 outer folds, mean *β* coefficients were calculated across 5 outer folds, and the median value across 100 permutations is plotted).

**Figure 5.**
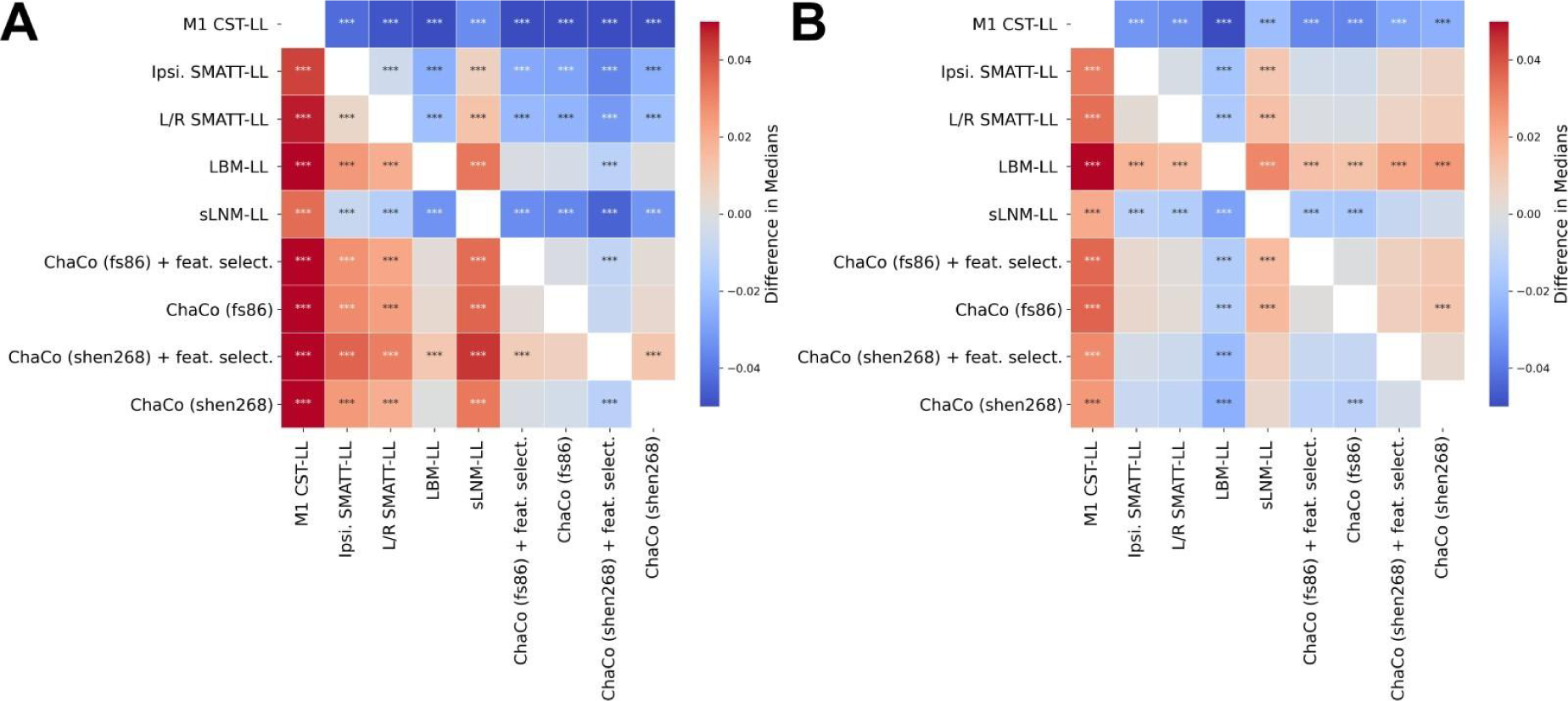
Statistical comparison of model performance for predicting motor scores using Mann-Whitney signed-rank tests. Colours shown indicate the differences in median explained variance scores for each model. **A.** Models trained using all (acute and chronic) training data. **B**. Models trained only using chronic data. *** denotes corrected *p <* 0.001 after Bonferroni correction. A positive difference indicates that the model on the y-axis (vertical) has a greater explained variance than the model on the *x*-axis (horizontal).

Within the theory-based biomarkers, M1-CST-LL models performed worse than ipsilesional SMATT-LL models (difference in *R*^2^ = -0.043, *p <* 0.001, 95% CI [*−*0.044*, −*0.041]) and worse than left/right SMATT-LL models (difference in *R*^2^ = -0.047, *p <* 0.001, 95% CI [*−*0.049*, −*0.045]).

Within the data-driven biomarkers, models using ChaCo scores parcellated with the Shen 268-region atlas and with correlation-based feature selection outperformed LBM-LL models (difference in *R*^2^ = -0.010, *p <* 0.001, 95% CI [*−*0.013*, −*0.007]). However, ChaCo models performed comparably to LBM-LL models when using only chronic training data (Supplementary Fig. 7A,B). Using all training data, all ChaCo models outperformed sLNM-LL models. When using only chronic training data, the differences between sLNM-LL models and ChaCo scores parcellated with the 268-region atlas were non-significant (Supplementary Fig. 7A,B). sLNM-LL models performed worse than LBM-LL models using all training data (difference in *R*^2^ = -0.034, *p <* 0.001, 95% CI [*−*0.035*, −*0.032]) and chronic training data (difference in *R*^2^ = -0.029, *p <* 0.001, 95% CI [*−*0.031*, −*0.028]).

For all models tested, ensemble models combining predictions from demographic data (subjects’ age, sex, and time since stroke) had better predictions than base models (Figure 6A-E, Table 2). Similarly, ensemble models merging predictions with the best-performing ChaCo models performed better than base lesion-load models. With the exception of LBM-LL models, ensemble models combining information from demographic data as well as ChaCo scores performed best (Figure 6A-E, Table 2). The best overall ensemble model included LBM-LL and 268-region ChaCo scores with feature selection.

**Figure 6.**
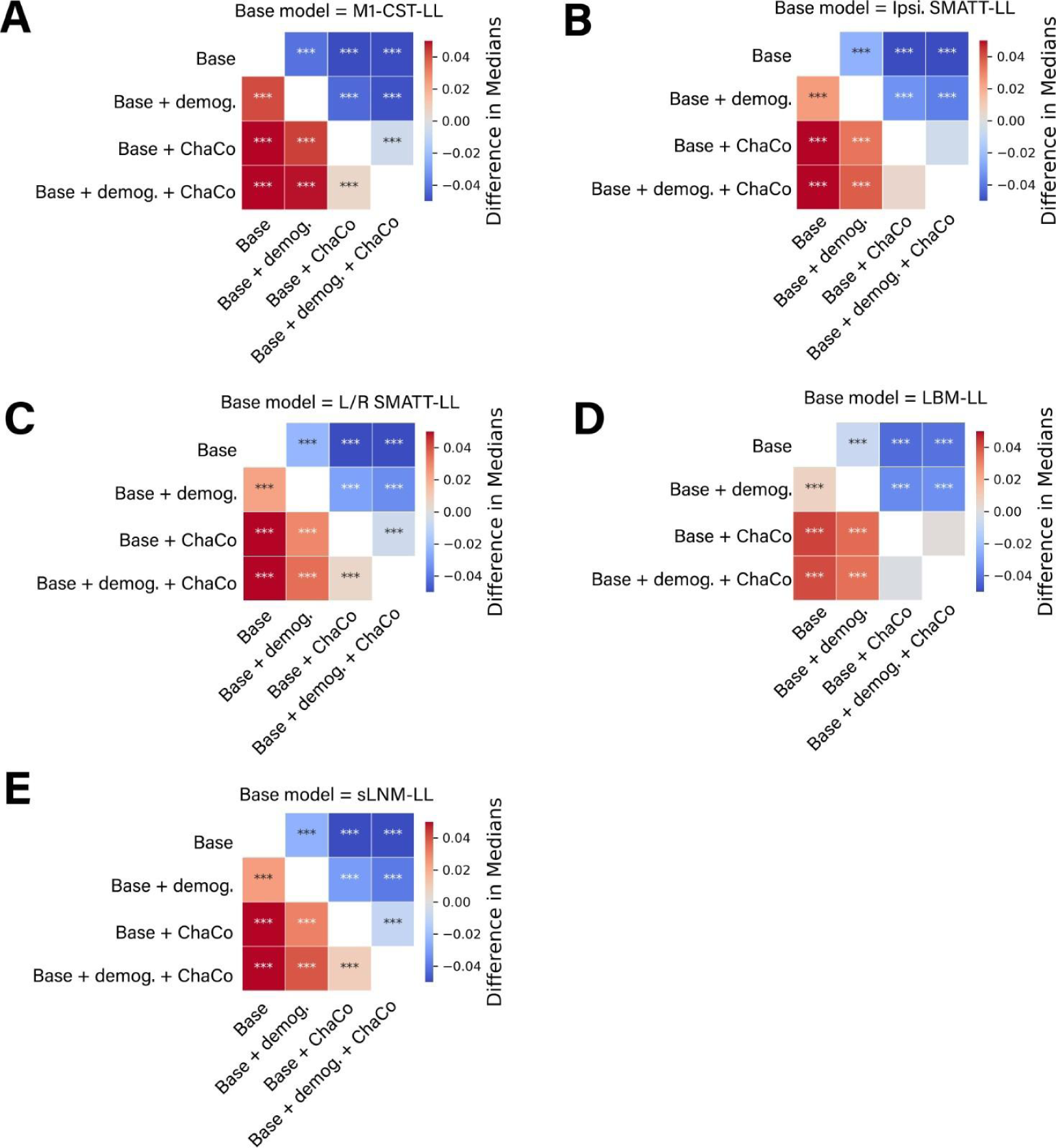
Statistical comparison of model performance for ensemble models. Demog. = demographic information (age, sex, days since stroke). ChaCo = model using 268-region ChaCo scores w/ feature selection. Significance of differences in explained variance were evaluated using Mann-Whitney signed-rank tests; ***denotes corrected *p <* 0.001 after Bonferroni correction. A positive difference value indicates that the model on the y-axis (vertical) has a greater explained variance than the model on the x-axis (horizontal).

**Table 2:**
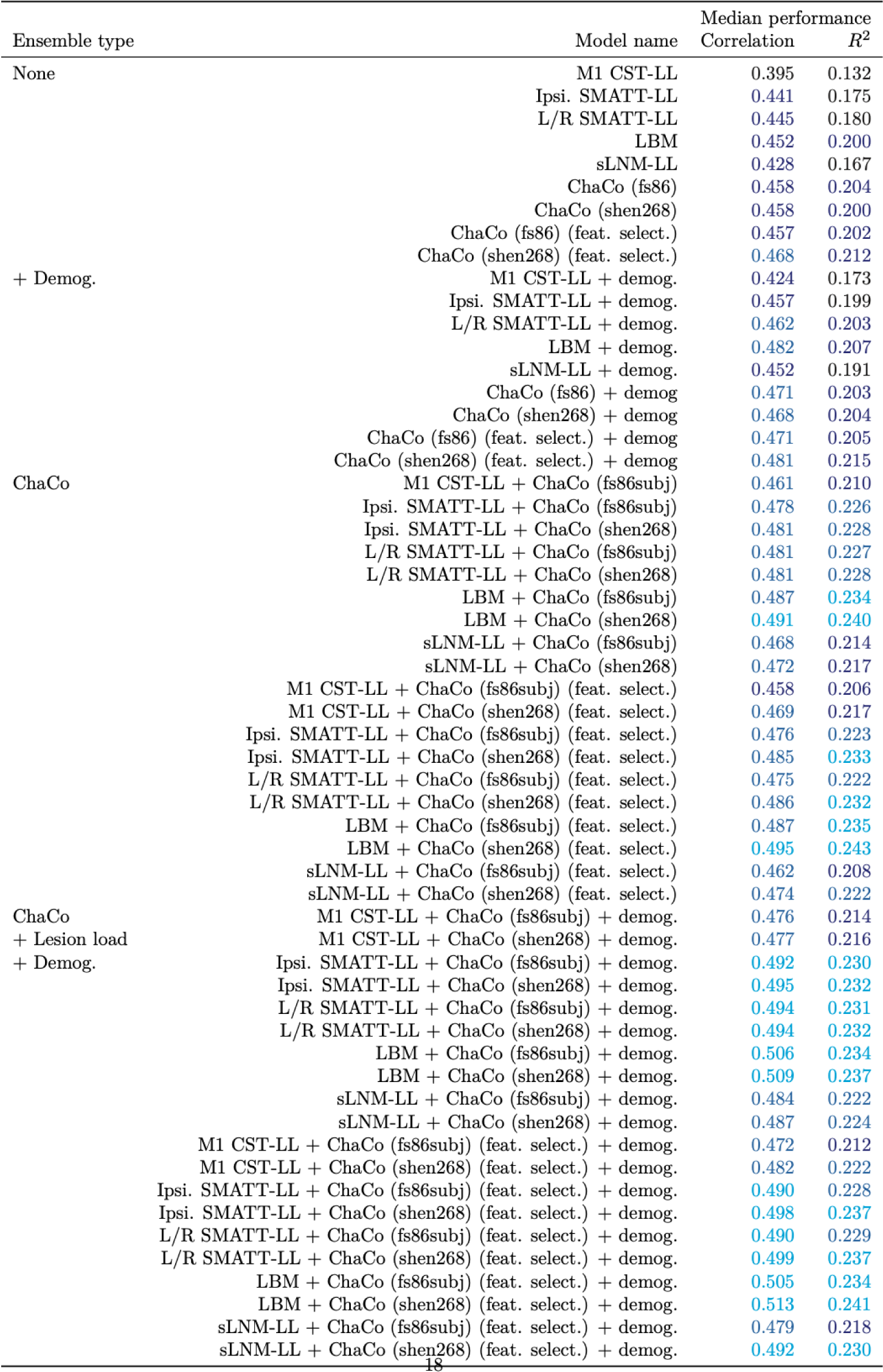
Test performance of all models evaluated, displaying median *R*^2^ and median correlation of average hold-out performances (i.e. average across 5 outer folds) across 100 permutations. Lighter blue shades represent better performance, and models are listed from theory-based to data-driven. Bemog. = demographics, Ipsi. = Ipsilesional, SMATT LL = sensorimotor tract template lesion load, L/R SMATT LL = left and right sensorimotor tract template lesion load, M1 CST LL = M1 corticospinal tract lesion load, ChaCo = Change in Connectivity, fs86 = FreeSurfer 86-region atlas, feat. select. = feature selection

### Featured selected by ChaCo models

Model weights for the best-performing ChaCo models are shown in Figure 4C, D, reflecting the median regression weight for each region across 100 train/test splits. There were several spatial similarities in the pattern of regression weights for the 86-region ChaCo model and 268-region ChaCo model with feature selection. For both atlases, negative model weights (indicating that more disconnection is associated with worse motor outcomes, holding all other factors constant) are assigned to left and right motor areas, as well as subcortical structures like the putamen and thalamus, whereas positive weights are assigned to frontal, parietal, and cingulate areas. In the 268-region ChaCo models, more regions in the right hemisphere are consistently included in the model than the left hemisphere.

The average correlation in feature weights between training folds was stable for the 268-region ChaCo score models, with an average r = 0.79 (Figure 7). Furthermore, many regions with high magnitude median feature weights had consistent weights across training folds. Finally, we observed evidence that 268-region ChaCo models were able to distinguish two regions’ relationships to motor outcomes, despite those regions being frequently damaged together (Supplementary Fig. 8).

**Figure 7.**
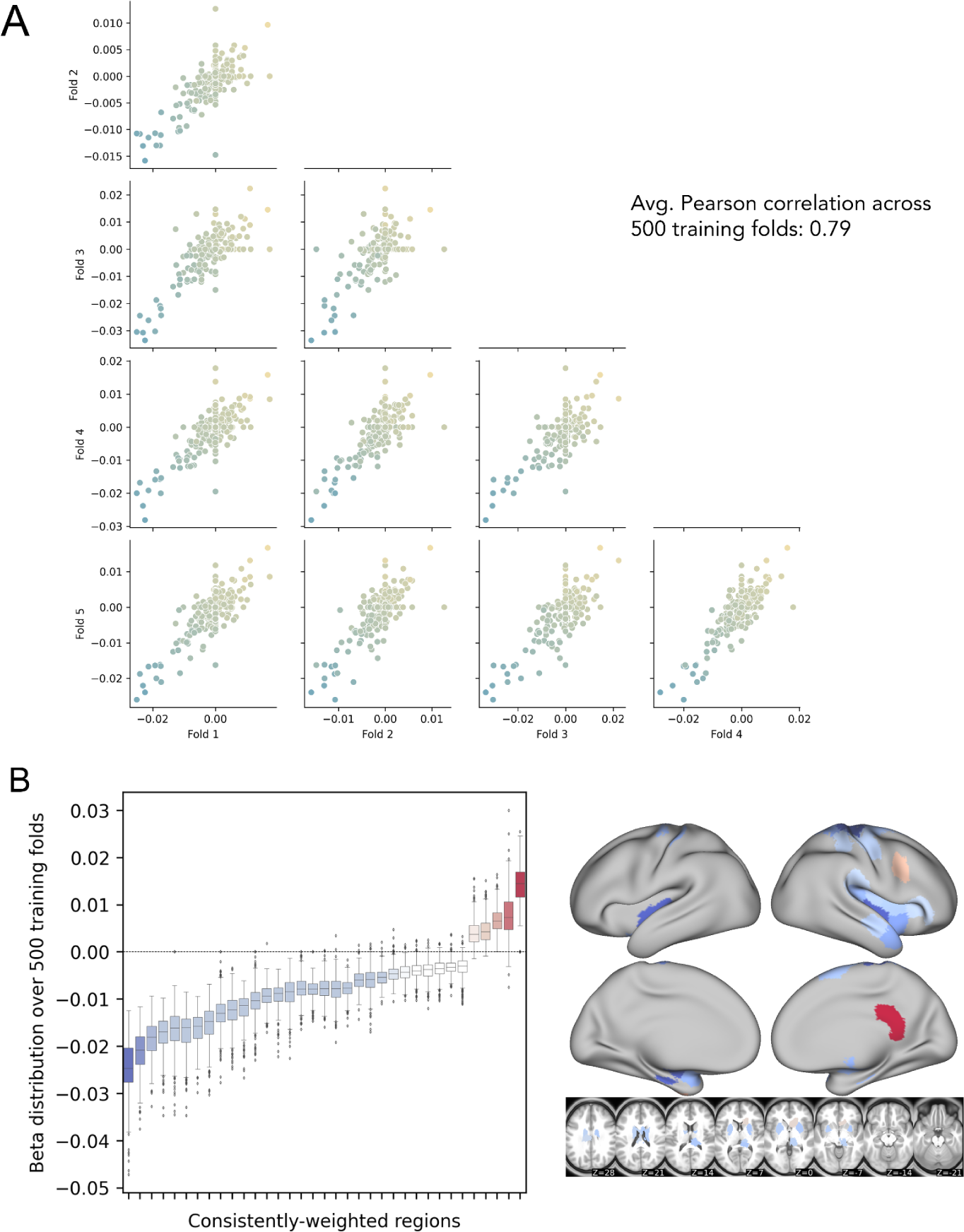
Analysis of feature stability for 268-region ChaCo models (with feature selection) and investigation of paradoxical feature weights. **A**. Correlation between beta coefficients across five training folds for one permutation. Each point corresponds to one region, and points are coloured by the mean beta coefficient for that region across 500 training folds (i.e. coloured based on y-axis value). **B.** Boxplots show the distribution of beta coefficients of consistently-weighted regions (defined as having median beta coefficients that are zero or of an opposite sign <5% of the time). In total, 30 regions with consistent negative weights and 5 regions with consistent positive weights remained. Median weights for consistently-weighted regions are plotted on a brain. The boxes extend from the lower to upper quartile values of the data, with a line at the median. Whiskers represent the range of the data from [Q1-1.5*IQR, Q3+1.5*IQR].

### Replication with subset data with FMA-UE scores

When predicting FMA-UE scores only, LBM-LL models performed best of all models tested (R^2^ = 0.145, p < 0.001), and performed significantly better than all theory-based models (Supplementary Fig. 9), though all models were still significant compared to chance.

## Discussion

In this study, we compared the performance of several structural imaging biomarkers in their ability to predict post-stroke motor outcomes. We found that, in general, data-driven models performed better than theory-based models in their ability to predict motor deficits in out-of-sample data, and this was replicated with a subset of the original data when predicting only FMA-UE scores. Among the data-driven models, we found that the best performance was obtained by modelling lesion damage using regional ChaCo scores. Contrary to our hypothesis, models using lesion-behaviour maps performed significantly better than structural lesion-network maps. Finally, we saw that combining predictions from demographic information and combining multiple biomarkers improved prediction of post-stroke motor ability over baseline models.

### Data-driven biomarkers outperform theory-based biomarkers

Using all training data, the best performing data-driven models used regional structural disconnection scores. These models, in addition to another data-driven biomarker, the extent of lesion damage to lesion-behaviour maps, outperformed all theory-based biomarkers.

Data-driven biomarkers may have outperformed theory-based biomarkers for two reasons. First, there may be regions outside of the primary motor system where lesions have an impact on motor performance. Damage to higher order motor areas in the frontal and parietal lobes^41^ that have been implicated in motor planning and execution,^42^ as well as damage to regions important for attention^43^ may be causally related to chronic motor outcomes. The same rationale underlies the most successful theory-driven models in a previous prediction study.^18^ Further, a patient’s ability to recover from or compensate for deficits may depend on a larger extent of lesion damage, the related overall stroke outcome, and related physiological consequences, such as autonomic dysfunction^44^ or inflammatory processes.^45^ Second, there may be features that are not causally related to motor function but are nonetheless predictive of long-term motor deficits. An *in silico* study has shown that imaging features with peak anatomo-clinical correlations can be located outside of the true neural correlates of a deficit^46^; in this specific example, damage in a temporal area correlated highly with a deficit that originated from either inferior parietal or inferior frontal damage. This can be explained by the typical lesion anatomy, which does not damage anatomical structures independently, but in highly systematic patterns imposed by the typical anatomy of the brain vasculature.^47^ Moreover, information from outside critical areas may supplement information in critical areas. For example, an imaging feature within a critical brain region or network may be damaged either by a small lacunar lesion that only causes a minor deficit which can be compensated for, or by a large lesion that fully disrupts a functional brain module and causes an irrevocable deficit. Damage to features outside of the critical area and that are indicative of a larger lesion might enable differentiation of these cases.

Hence, the direct damage to specific motor structures alone may not be optimally predictive of poor outcomes. In this paper, structural disconnection of areas outside of the primary motor system was predictive of worse motor outcomes. Similarly, the extent of damage to a lesion behaviour map including voxels that lie mostly outside of the motor system (Supplementary Fig. 10) was able to predict motor outcomes better than damage to known motor tracts. This study suggests that regardless of whether these extra-primary motor structures are causally related to a deficit, they are more useful biomarkers of chronic motor deficits than the extent of damage to white matter tracts of the motor system that are currently implemented in prediction studies.

### Lesion-based structural disconnectivity for prediction

Models using ChaCo scores performed best of all models tested, particularly when feature selection was employed. These are high-dimensional models that may require more data to start outperforming simpler models,^22^ which may explain the drop in their relative performance when using smaller subsets of the data for training, including using only chronic data and using only subjects with FMA-UE scores. However, with sufficient data, one strength of models using ChaCo scores can be understood, in part, by considering how lesion data is represented relative to LBM and sLNM models. For LBM and sLNM, the data on which feature selection takes place are voxels. On the other hand, in ChaCo models, the data on which feature selection takes place are regional measures of structural disconnection. This data transformation essentially reduces the number of “rare” features compared to voxelwise representations (Supplementary Fig. 11). Statistical power may be increased when voxelwise lesion data is first transformed into regional ChaCo scores; non-overlapping lesions that affect different portions of the same tract are mapped onto ChaCo scores of the same region or set of regions. The drawback to this approach is that regional ChaCo scores do not enable the detection of associations between damage to specific tracts and motor outcomes; if such associations exist then that signal may be diluted in regional measures.

### Feature weights of the ChaCo models

In this paper, our main aim was to identify stroke imaging biomarkers that predict motor outcome. We did not aim to uncover the precise nature of the neural correlates of motor deficits. However, further investigation of model features and their weights can provide some clarity in understanding the model’s performance.

Several grey matter regions that are part of the known motor system were incorporated into ChaCo models with negative weights, suggesting that more damage to these regions is associated with worse motor outcomes. Such regions include the primary somatomotor cortex and subcortical structures, as well as secondary motor structures in frontal and parietal cortices. Many regions that were consistently assigned negative weights were neighbouring regions, in line with the spatial distribution of motor networks and somatotopy of the motor system. However, several regions, in particular in the right frontal cortex and medial surface, were consistently assigned a positive weight. In other words, some brain regions existed for which feature weights indicated a paradoxical lesion-deficit relationship in the sense that brain damage was linked to a more favourable motor outcome. Some cases of genuine facilitation due to brain damage have been documented,^24,48^ and inhibitory interregional brain modes that can explain paradoxical lesion effects are assumed.^49,50^ Hence, paradoxical lesion effects underlying motor outcome may provide a counter-intuitive, but still viable explanation of our findings. On the other hand, methodological aspects could also be an explanation for apparently paradoxical effects. First, paradoxical associations might arise as an artefact from the lesion anatomy.^51^ For illustration, imagine a stroke population in which some patients suffer from visual field defects after posterior brain damage to the visual cortex or the optic radiation. The existence of a frontal lesion might then be anticorrelated with visual field defects - not because of a true paradoxical lesion effect due to inhibition, but as a mere statistical effect following from the lesion anatomy: a patient with a frontal lesion is unlikely to simultaneously suffer from a posterior lesion and, hence, is unlikely to suffer from visual field defects. Similar effects are imaginable on a smaller scale affecting neighbouring brain regions.^51^

Second, paradoxical effects might also emerge as a simple statistical artefact. The feature weights in a high-dimensional prediction model can be unstable^33^ and, especially with highly correlated data, can somehow be decoupled from causality and the actual structure of the investigated entity. In our study, the stability of features was decent, though still markedly inferior to some previous studies that explicitly optimised feature replicability to create interpretable high-dimensional models.^52,53^ Accordingly, we identified some areas where paradoxical feature weights emerged but were unstable across replications in subsamples. Only for some areas, the paradoxical feature weights were stable across replications. In the current study, we are unable to come to final conclusions regarding the generalizability of these paradoxical weights. Future studies are needed to validate or optimise our modelling and model interpretation strategies.

### Shared variance dimensionality reduction approaches: diluting signals that should be separated?

Structural disconnection-behaviour mapping has been recently employed to identify white matter correlates of motor function,^54^ and to predict 2-week motor scores from lesion-induced pairwise structural disconnection.^7^ In the latter, most conceptually similar to this study, tractography is seeded from voxels in a lesion, and a map representing the probability of disconnection from the lesion is generated. Features for ridge regression models are generated via principal components analysis (PCA) on the voxelwise structural disconnectivity maps. This is a reasonable dimensionality reduction step that has been applied in voxelwise lesion-symptom mapping to identify primary axes of variance, but is a fundamentally different approach for representing structural disconnection compared to regional ChaCo scores derived from NeMo tool, in which dimensionality is reduced by summarising the degree of disconnection for each brain region separately. As demonstrated in this paper, regions that are frequently damaged together (and whose tracts may load onto the same principal component in the PCA approach) may have opposite relationships to motor scores, so dimensionality reduction steps based on shared variance that do not consider relationships between structural disconnections and outcome variables may compress relevant individual signals into single features, reducing the predictive performance of a model.

### Surprisingly strong performance of a simple biomarker: LBM lesion load

We hypothesised that because of previously-identified relationships between structural disconnections and motor deficits, lesion load to structural lesion-network maps would outperform lesion load to basic lesion-behaviour maps. On the contrary, we saw that lesion load to a lesion-behaviour map performed better than structural lesion network maps. In some cases, lesion-behaviour map lesion-load performed as well as complex, high-dimensional ChaCo models. This lesion-behaviour map was derived from an independent dataset, suggesting that the voxelwise feature selection performed produced a map of association that is generalizable to new data. Predicting motor deficits using lesion load to a lesion-behaviour map can be done with simple linear regression, making this biomarker accessible to those with a limited coding background. Hence, even though a single lesion load measure might at first glance appear to be overly simple and unfit to represent the complexity of the human brain and its pathology, it might still provide a biomarker that can be meaningful in clinical studies with simple, straightforward interpretable design. However, high-precision personalised medicine should rely on more complex, high dimensional imaging markers such as ChaCo disconnection scores.

### Higher-order motor areas are relevant for motor outcome prediction

This study supports previous results showing that M1-CST-LL alone may not capture variance associated with chronic post-stroke motor outcomes as well as other metrics,^16,17,19^ although few studies have directly compared the out-of-sample performance of M1-CST-LL against more complex metrics like those assessed in this paper. Damage to descending fibres from higher order motor areas has been associated with motor deficits after stroke,^17,55^ and may explain additional variance in chronic motor outcome that is not captured by damage to the M1 corticospinal tract alone. This study supports the notion that further development of non-M1-CST biomarkers is justified and that taking into account damage to non-primary motor regions may be important for understanding variation in motor outcome.

### Ensemble models

Finally, we saw that averaging predictions from multiple models generally improves performance. This suggests that the information captured by each data type is not redundant, and that using multiple different lesion metrics may compensate for weaknesses of different feature representations. In this study, SMATT-LL models have the ability to distinguish between damage to several adjacent descending corticospinal tracts, but are unable to measure lesion-deficit associations outside this area, or in any grey matter regions. On the other hand, ChaCo models can detect associations across the entire brain and in the grey matter but may not have sufficient resolution between different descending motor tracts to capture relevant variance there. Beyond predictions of chronic motor scores, prediction models may be improved by testing and possibly combining multiple features as well as multiple feature representations (specifically, LBM-LL and ChaCo scores) to obtain an optimal model.^21,56^

### Limitations

There are several limitations of this study. Without baseline motor scores, we cannot evaluate the relative predictive power of lesion damage versus baseline behavioural information, which may share variance. Since this information would likely be accessible for future clinical trials or prediction contexts, it is important to determine to what degree lesion information provides explanatory power above baseline motor scores. Previous studies have shown that prediction models with behavioural predictors and imaging biomarkers explain more variance in motor outcomes than the models with behavioural predictors alone^14^ Similarly, the lack of subject-level rehabilitation data is another limitation. Patients likely completed different levels of rehabilitation, which would impact their chronic motor scores. Ultimately, any additional variance that is not captured by the model limits the predictive accuracy of the models and future predictive work should attempt to incorporate as many recovery-relevant variables as possible. Additionally, the inclusion of metrics that are not specific to motor deficits (i.e., NIHSS) may have reduced the performance of models.

Finally, the strength of LBM-LL/sLNM-LL models relative to ChaCo models may be reduced because of the distribution shift in the training vs. testing dataset: the sample used to generate the LBM was different from the sample on which it was tested, whereas for ChaCo models, feature selection was performed using the same dataset on which the models were tested.

## Acknowledgements

We thank the feedback from all members of the ENIGMA consortium, and for the patients who made their data available for secondary analyses.

## Supplementary material

Supplementary figures can be found in the supplementary material.

## References

1. Kelly-Hayes M, Beiser A, Kase CS, Scaramucci A, D’Agostino RB, Wolf PA. The influence of gender and age on disability following ischemic stroke: the Framingham study. J Stroke Cerebrovasc Dis. 2003;12(3):119–126.

2. Bonkhoff AK, Grefkes C. Precision medicine in stroke: towards personalized outcome predictions using artificial intelligence. Brain. 2022;145(2):457–475.

3. Boyd LA, Hayward KS, Ward NS, et al. Biomarkers of stroke recovery: Consensus-based core recommendations from the Stroke Recovery and Rehabilitation Roundtable. Int J Stroke. 2017;12(5):480–493.

4. Tozlu C, Edwards D, Boes A, et al. Machine Learning Methods Predict Individual Upper-Limb Motor Impairment Following Therapy in Chronic Stroke. Neurorehabil Neural Repair. 2020;34(5):428–439.

5. Kuceyeski A, Navi BB, Kamel H, et al. Structural connectome disruption at baseline predicts 6-months post-stroke outcome. Hum Brain Mapp. 2016;37(7):2587–2601.

6. Griffis JC, Metcalf NV, Corbetta M, Shulman GL. Structural Disconnections Explain Brain Network Dysfunction after Stroke. Cell Rep. 2019;28(10):2527–2540.e9.

7. Salvalaggio A, De Filippo De Grazia M, Zorzi M, Thiebaut de Schotten M, Corbetta M. Post-stroke deficit prediction from lesion and indirect structural and functional disconnection. Brain. Published online June 23, 2020. doi:10.1093/brain/awaa156

8. Bowren M, Bruss J, Manzel K, et al. Post-stroke outcomes predicted from multivariate lesion-behaviour and lesion network mapping. Brain. Published online January 13, 2022. doi:10.1093/brain/awac010

9. Zhu LL, Lindenberg R, Alexander MP, Schlaug G. Lesion load of the corticospinal tract predicts motor impairment in chronic stroke. Stroke. 2010;41(5):910–915.

10. Feng W, Wang J, Chhatbar PY, et al. Corticospinal tract lesion load: An imaging biomarker for stroke motor outcomes. Ann Neurol. 2015;78(6):860–870.

11. Findlater SE, Hawe RL, Mazerolle EL, et al. Comparing CST Lesion Metrics as Biomarkers for Recovery of Motor and Proprioceptive Impairments After Stroke. Neurorehabil Neural Repair. 2019;33(10):848–861.

12. Lam TK, Dawson DR, Honjo K, et al. Neural coupling between contralesional motor and frontoparietal networks correlates with motor ability in individuals with chronic stroke. J Neurol Sci. 2018;384:21–29.

13. Pineiro R, Pendlebury ST, Smith S, et al. Relating MRI changes to motor deficit after ischemic stroke by segmentation of functional motor pathways. Stroke. 2000;31(3):672–679.

14. Kim B, Winstein C. Can Neurological Biomarkers of Brain Impairment Be Used to Predict Poststroke Motor Recovery? A Systematic Review. Neurorehabil Neural Repair. 2017;31(1):3–24.

15. Paul T, Cieslak M, Hensel L, et al. The role of corticospinal and extrapyramidal pathways in motor impairment after stroke. Brain Commun. 2023;5(1):fcac301.

16. Park CH, Kou N, Ward NS. The contribution of lesion location to upper limb deficit after stroke. J Neurol Neurosurg Psychiatry. 2016;87(12):1283–1286.

17. Ito KL, Kim B, Liu J, et al. Corticospinal Tract Lesion Load Originating From Both Ventral Premotor and Primary Motor Cortices Are Associated With Post-stroke Motor Severity. Neurorehabil Neural Repair. 2022;36(3):179–182.

18. Rondina JM, Filippone M, Girolami M, Ward NS. Decoding post-stroke motor function from structural brain imaging. Neuroimage Clin. 2016;12:372–380.

19. Rondina JM, Park CH, Ward NS. Brain regions important for recovery after severe post-stroke upper limb paresis. J Neurol Neurosurg Psychiatry. 2017;88(9):737–743.

20. Schulz R, Park CH, Boudrias MH, Gerloff C, Hummel FC, Ward NS. Assessing the integrity of corticospinal pathways from primary and secondary cortical motor areas after stroke. Stroke. 2012;43(8):2248–2251.

21. Kasties V, Karnath HO, Sperber C. Strategies for feature extraction from structural brain imaging in lesion-deficit modelling. Hum Brain Mapp. 2021;42(16):5409–5422.

22. Bourached A, Bonkhoff AK, Schirmer MD, et al. Scaling behaviors of deep learning and linear algorithms for the prediction of stroke severity. bioRxiv. Published online December 6, 2022. doi:10.1101/2022.12.05.22283102

23. Bzdok D, Engemann D, Thirion B. Inference and Prediction Diverge in Biomedicine. Patterns (N Y). 2020;1(8):100119.

24. Sperber C, Griffis J, Kasties V. Indirect structural disconnection-symptom mapping. Brain Struct Funct. Published online September 1, 2022. doi:10.1007/s00429-022-02559-x

25. Kuceyeski A, Maruta J, Relkin N, Raj A. The Network Modification (NeMo) Tool: elucidating the effect of white matter integrity changes on cortical and subcortical structural connectivity. Brain Connect. 2013;3(5):451–463.

26. Cramer SC. Stratifying Patients With Stroke in Trials That Target Brain Repair. Stroke. 2010;41(10_suppl_1):S114-S116.

27. Richards LG, Cramer SC. Advances in Stroke Recovery Therapeutics. Stroke. 2022;53(1):260–263.

28. Liew SL, Zavaliangos-Petropulu A, Jahanshad N, et al. The ENIGMA Stroke Recovery Working Group: Big data neuroimaging to study brain-behavior relationships after stroke. Hum Brain Mapp. Published online April 20, 2020. doi:10.1002/hbm.25015

29. Gladstone DJ, Danells CJ, Black SE. The fugl-meyer assessment of motor recovery after stroke: a critical review of its measurement properties. Neurorehabil Neural Repair. 2002;16(3):232–240.

30. Sulter G, Steen C, De Keyser J. Use of the Barthel index and modified Rankin scale in acute stroke trials. Stroke. 1999;30(8):1538–1541.

31. Lyden P. Using the National Institutes of Health Stroke Scale: A Cautionary Tale. Stroke. 2017;48(2):513–519.

32. Hastie T, Friedman J, Tibshirani R. The Elements of Statistical Learning. Springer New York; 2001.

33. Tian Y, Zalesky A. Machine learning prediction of cognition from functional connectivity: Are feature weights reliable? Neuroimage. 2021;245:118648.

34. Archer DB, Vaillancourt DE, Coombes SA. A Template and Probabilistic Atlas of the Human Sensorimotor Tracts using Diffusion MRI. Cereb Cortex. 2018;28(5):1685–1699.

35. Pustina D, Avants B, Faseyitan OK, Medaglia JD, Coslett HB. Improved accuracy of lesion to symptom mapping with multivariate sparse canonical correlations. Neuropsychologia. 2018;115:154–166.

36. Desikan RS, Ségonne F, Fischl B, et al. An automated labeling system for subdividing the human cerebral cortex on MRI scans into gyral based regions of interest. Neuroimage. 2006;31(3):968–980.

37. Fischl B, Salat DH, Busa E, et al. Whole brain segmentation: automated labeling of neuroanatomical structures in the human brain. Neuron. 2002;33(3):341–355.

38. Shen X, Tokoglu F, Papademetris X, Constable RT. Groupwise whole-brain parcellation from resting-state fMRI data for network node identification. Neuroimage. 2013;82:403–415.

39. Guyon I, Elisseeff A. An introduction to variable and feature selection. J Mach Learn Res. 2003;3:1157–1182.

40. Tibshirani R. Regression Shrinkage and Selection via the Lasso. J R Stat Soc Series B Stat Methodol. 1996;58(1):267–288.

41. Galea MP, Darian-Smith I. Multiple corticospinal neuron populations in the macaque monkey are specified by their unique cortical origins, spinal terminations, and connections. Cereb Cortex. 1994;4(2):166–194.

42. Ball T, Schreiber A, Feige B, Wagner M, Lücking CH, Kristeva-Feige R. The role of higher-order motor areas in voluntary movement as revealed by high-resolution EEG and fMRI. Neuroimage. 1999;10(6):682–694.

43. Rinne P, Hassan M, Fernandes C, et al. Motor dexterity and strength depend upon integrity of the attention-control system. Proc Natl Acad Sci U S A. 2018;115(3):E536–E545.

44. Xiong L, Tian G, Leung H, et al. Autonomic Dysfunction Predicts Clinical Outcomes After Acute Ischemic Stroke: A Prospective Observational Study. Stroke. 2018;49(1):215–218.

45. Shi K, Tian DC, Li ZG, Ducruet AF, Lawton MT, Shi FD. Global brain inflammation in stroke. Lancet Neurol. 2019;18(11):1058–1066.

46. Mah YH, Husain M, Rees G, Nachev P. Human brain lesion-deficit inference remapped. Brain. 2014;137(Pt 9):2522–2531.

47. Zhao Y, Halai AD, Lambon Ralph MA. Evaluating the granularity and statistical structure of lesions and behaviour in post-stroke aphasia. Brain Commun. 2020;2(2):fcaa062.

48. Kapur N. Paradoxical functional facilitation in brain-behaviour research. A critical review. Brain. 1996;119 (Pt 5):1775–1790.

49. Valero-Cabré A, Toba MN, Hilgetag CC, Rushmore RJ. Perturbation-driven paradoxical facilitation of visuo-spatial function: Revisiting the “Sprague effect.” Cortex. 2020;122:10–39.

50. Toba MN, Godefroy O, Rushmore RJ, et al. Revisiting “brain modes” in a new computational era: approaches for the characterization of brain-behavioural associations. Brain. 2020;143(4):1088–1098.

51. Sperber C, Karnath HO. Inhibition between human brain areas or methodological artefact? Brain. 2020;143(5):e38–e38.

52. Zhang Y, Kimberg DY, Coslett HB, Schwartz MF, Wang Z. Multivariate lesion-symptom mapping using support vector regression. Hum Brain Mapp. 2014;35(12):5861–5876.

53. Wiesen D, Sperber C, Yourganov G, Rorden C, Karnath HO. Using machine learning-based lesion behavior mapping to identify anatomical networks of cognitive dysfunction: Spatial neglect and attention. Neuroimage. 2019;201:116000.

54. Wawrzyniak M, Stockert A, Klingbeil J, Saur D. Voxelwise structural disconnection mapping: Methodological validation and recommendations. Neuroimage Clin. 2022;35:103132.

55. Riley JD, Le V, Der-Yeghiaian L, et al. Anatomy of stroke injury predicts gains from therapy. Stroke. 2011;42(2):421–426.

56. Park CH, Ohn SH. The predictive value of lesion and disconnectome loads for upper limb motor impairment after stroke. Neurol Sci. 2022;43(5):3097–3104.

