## Supplementary Materials for "Data-driven biomarkers outperform theory-based biomarkers in predicting stroke motor outcomes"

Supplementary Table 1: Number of subjects with data for each type of assessment, broken down by chronicity (acute-subacute/chronic) and site. FMA UE = Fugl Meyer Assessment of Upper Extremity, Barthel = Barthel Index, NIHSS = National Institutes of Health Stroke Score.

| **Acute/subacute** | | | | |
|---|---|---|---|---|
| **Site ID** | **Fugl Meyer UE** | **Barthel** | **NIHSS** | **Grip Strength** |
| r005 | 1 | 0 | 0 | 0 |
| r009 | 0 | 0 | 49 | 0 |
| r025 | 0 | 0 | 0 | 9 |
| r028 | 1 | 0 | 0 | 0 |
| r031 | 36 | 0 | 0 | 0 |
| r038 | 0 | 72 | 0 | 0 |
| r040 | 0 | 57 | 0 | 0 |
| r047 | 2 | 0 | 0 | 0 |
| r049 | 0 | 0 | 21 | 0 |
| r050 | 0 | 0 | 14 | 0 |
| r053 | 0 | 0 | 52 | 0 |
| r054 | 12 | 0 | 0 | 0 |
| **Chronic** | | | | |
| **Site ID** | **Fugl Meyer UE** | **Barthel** | **NIHSS** | **Grip Strength** |
| r001 | 39 | 0 | 0 | 0 |
| r002 | 12 | 0 | 0 | 0 |
| r003 | 15 | 0 | 0 | 0 |
| r004 | 19 | 0 | 0 | 0 |
| r005 | 27 | 0 | 0 | 0 |
| r009 | 60 | 0 | 0 | 0 |
| r025 | 0 | 0 | 0 | 16 |
| r027 | 28 | 0 | 0 | 0 |
| r028 | 21 | 0 | 0 | 0 |
| r031 | 1 | 0 | 0 | 0 |

| **Acute/subacute** | | | | |
|---|---|---|---|---|
| r034 | 15 | 0 | 0 | 0 |
| r035 | 15 | 0 | 0 | 0 |
| r038 | 0 | 18 | 0 | 0 |
| r040 | 0 | 14 | 0 | 0 |
| r042 | 22 | 0 | 0 | 0 |
| r044 | 4 | 0 | 0 | 0 |
| r045 | 4 | 0 | 0 | 0 |
| r046 | 11 | 0 | 0 | 0 |
| r047 | 44 | 0 | 0 | 0 |
| r048 | 43 | 0 | 0 | 0 |
| r052 | 0 | 0 | 32 | 0 |
| r053 | 0 | 0 | 2 | 0 |

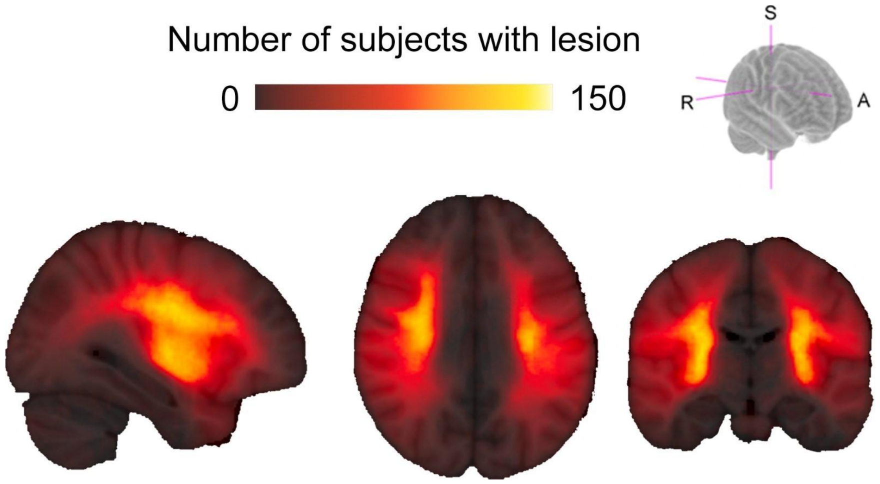


Supplementary Figure 1: Distribution of lesions in the ENIGMA cohort.

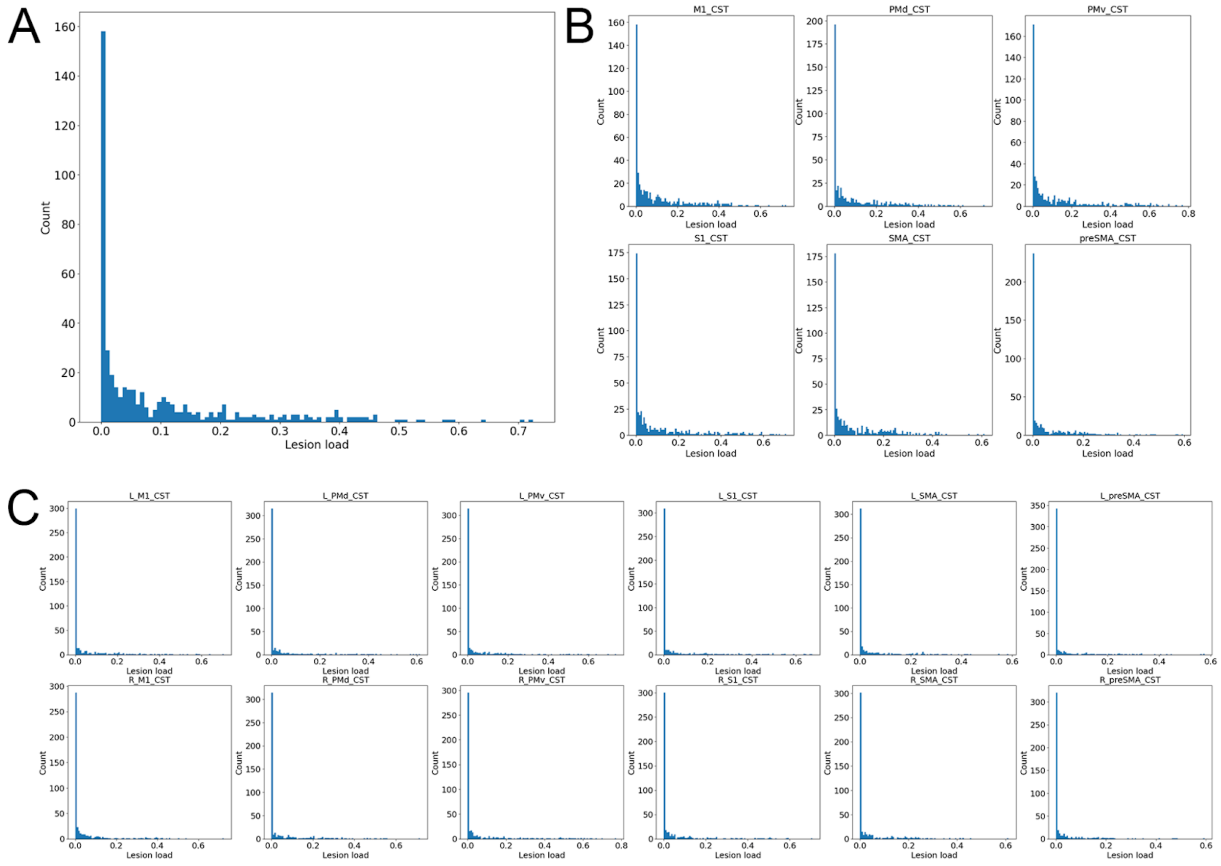


Supplementary Figure 2: Distribution of lesion load variables for chronic subjects. **A**. M1-CST-LL distribution, **B.** Ipsilesional SMATT-LL distribution, **C.** Bilateral SMATT-LL distribution.

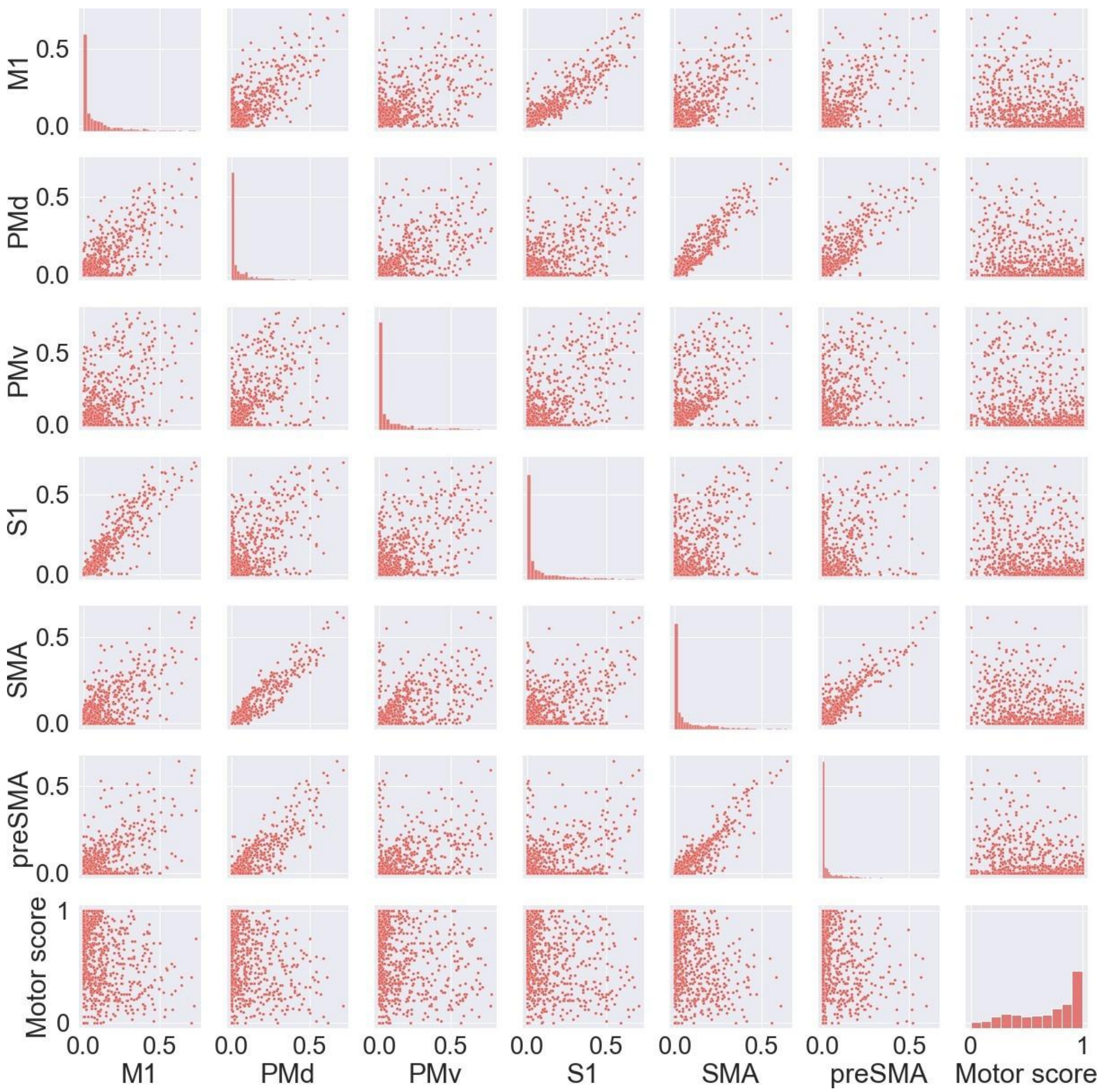


Supplementary Figure 3: Correlations between lesion load calculated for each ipsilesional tract in the sensorimotor area tract template atlas.

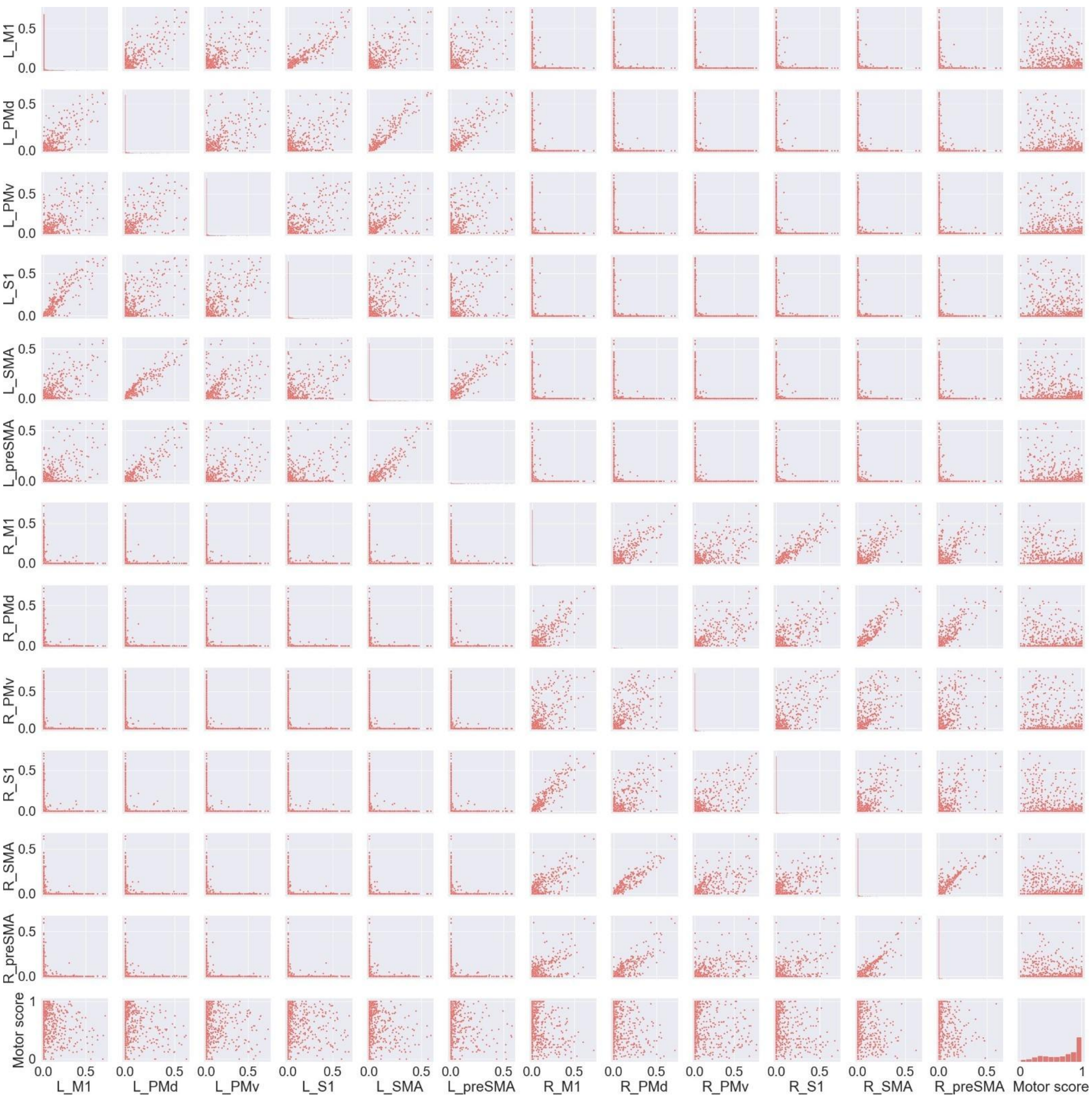


Supplementary Figure 4: Correlations between lesion load calculated for each left and right hemisphere tract in the sensorimotor area tract template atlas.

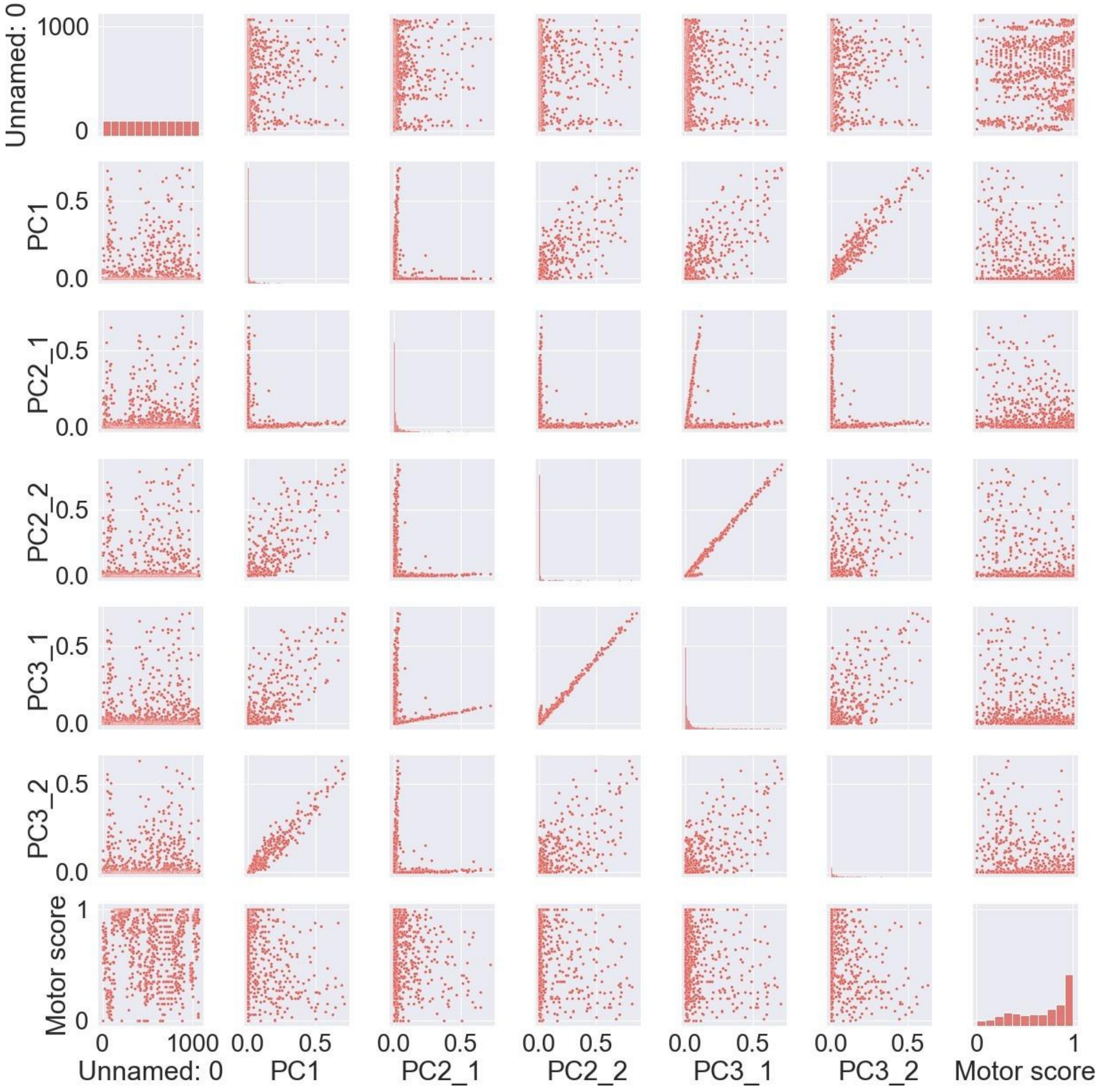


Supplementary Figure 5: Correlations between lesion load calculated for each structural lesion network-derived principal component map.

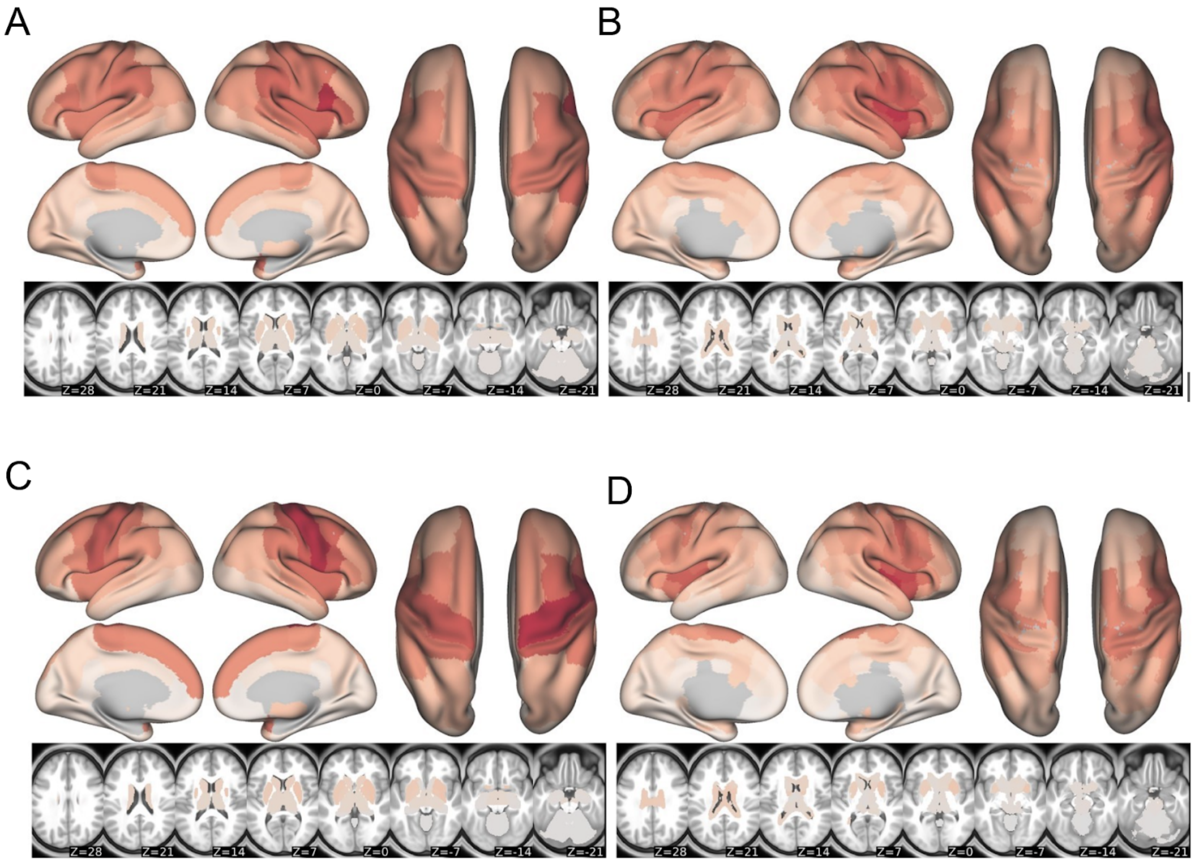


Supplementary Figure 6. Top: Mean regional change in connectivity (ChaCo) scores across acute and chronic subjects used in predictive models. Mean of ChaCo scores parcellated with (**A**) the FreeSurfer 86-region atlas (min. = 0.005, max. = 0.120) and with (**B**) the Shen 268-region atlas (min. = 0.0025, max. = 0.146), normalized to the maximum value across regions (red = higher mean ChaCo score across subjects) Standard deviation of regional change in connectivity (ChaCo) scores across acute and chronic subjects used in predictive models. Standard deviation of ChaCo scores parcellated with (**C**) the FreeSurfer 86-region atlas (min. = 0.024, max. = 0.26) and with (**D**) the Shen 268-region atlas (min. = 0.014, max. = 0.29), normalized to the maximum value across regions (red = higher standard deviation of ChaCo score across subjects)

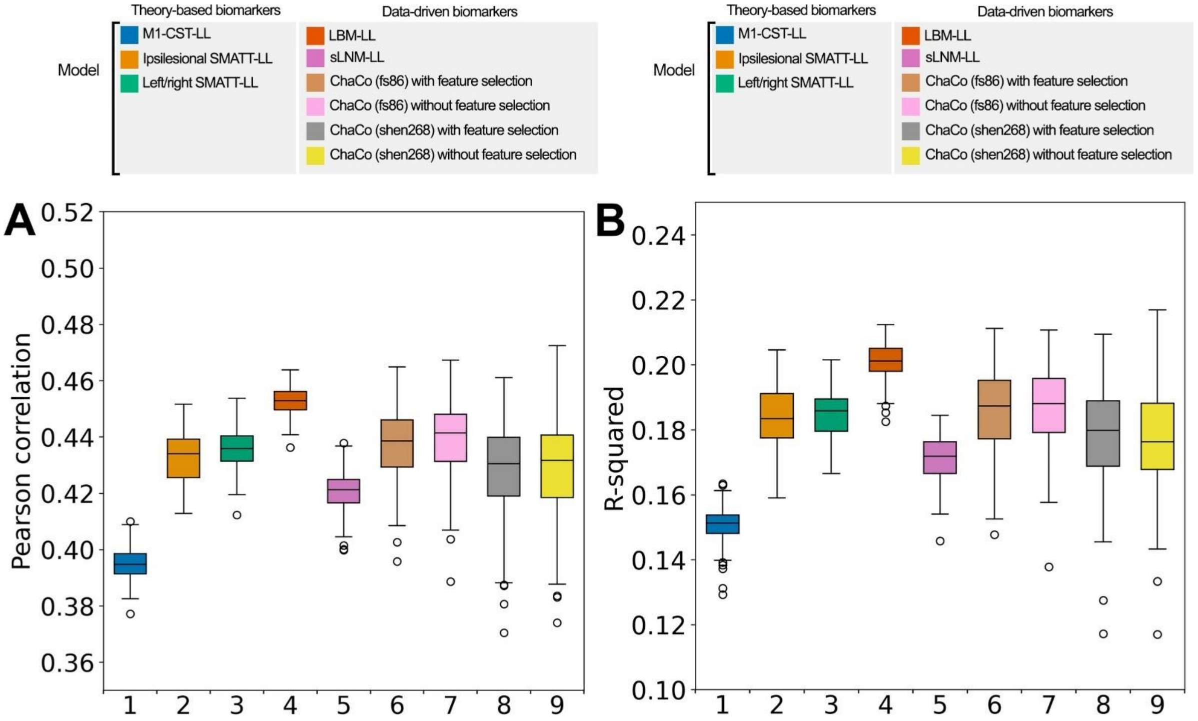


Supplementary Figure 7: Summary of model performance metrics across all models tested using only chronic data for training. **A.** and **B.** Distribution of model performance (mean Pearson correlation/$R^2$ across 5 outer folds for 100 permutations of the data). Boxplots are colored arbitrarily for clarity. The boxes extend from the lower to upper quartile values of the data, with a line at the median. Whiskers represent the range of the data from [Q1-1.5*IQR, Q3+1.5*IQR]

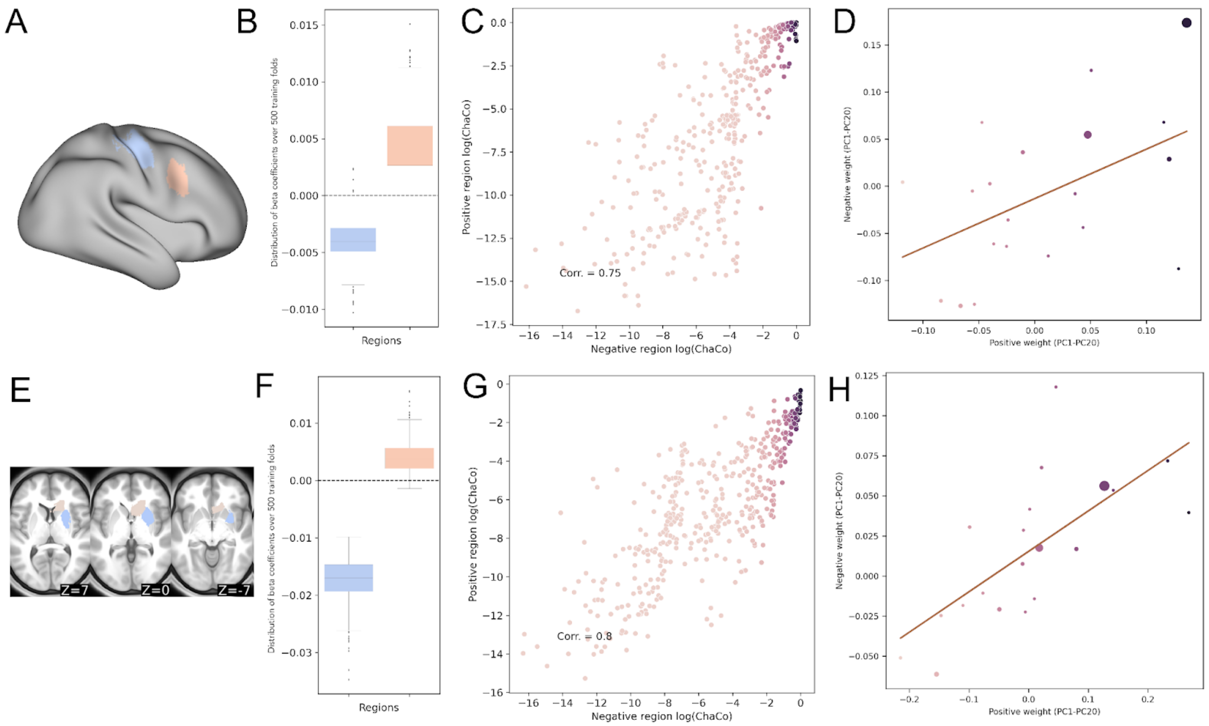


Supplementary Figure 8: The correlation between ChaCo scores for all pairs of consistently-weighted regions was calculated. In total, 4 pairs of regions had ChaCo scores with correlation > 0.8. Two such pairs are plotted in **A** and **E**, coloured based on the median beta coefficient for that region. **B, F.** distribution of beta coefficients for region-pairs with >0.8 correlation in ChaCo scores but opposite sign beta coefficients. **C, G.** Scatterplot showing distribution of ChaCo scores for each region-pair. **D, H.** Region-pairs' loading onto principal components that explain >90% of the variance in 268-region ChaCo score, with the negatively-weighted region on the y-axis and the positively-weighted region on the x-axis. Dots indicate the regions' loading onto each component, and the size of the dots corresponds to the percentage of variance explained in the full dataset by each component.

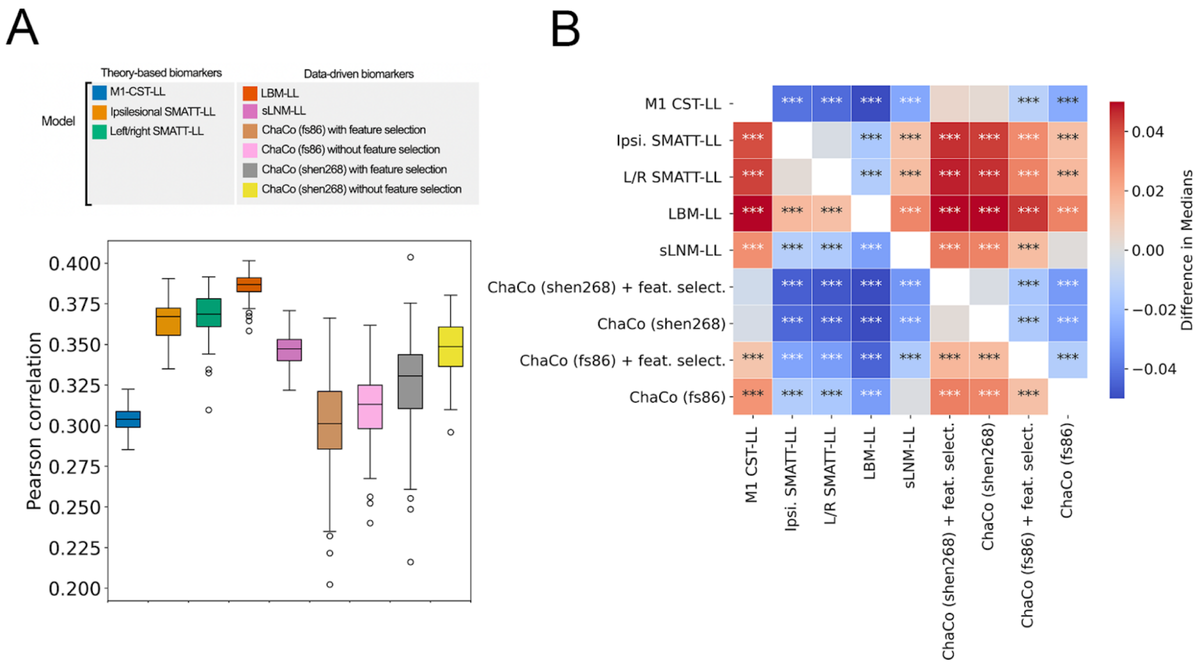


Supplementary Figure 9. Summary of model performance metrics across all models tested using only data with Fugl-Meyer motor scores (chronic and acute/subacute data). **A.** Distribution of model performance (mean Pearson correlation across 5 outer folds for 100 permutations of the data). Boxplots are coloured arbitrarily for clarity. The boxes extend from the lower to upper quartile values of the data, with a line at the median. Whiskers represent the range of the data from [Q1-1.5*IQR, Q3+1.5*IQR]. **B.** Statistical comparison of model performance for predicting motor scores using Mann-Whitney signed-rank tests. Colours shown indicate the differences in median explained variance scores for each model. *** denotes corrected $p < 0.001$ after Bonferroni correction. A positive difference indicates that the model on the y-axis (vertical) has a greater explained variance than the model on the *x*-axis (horizontal).

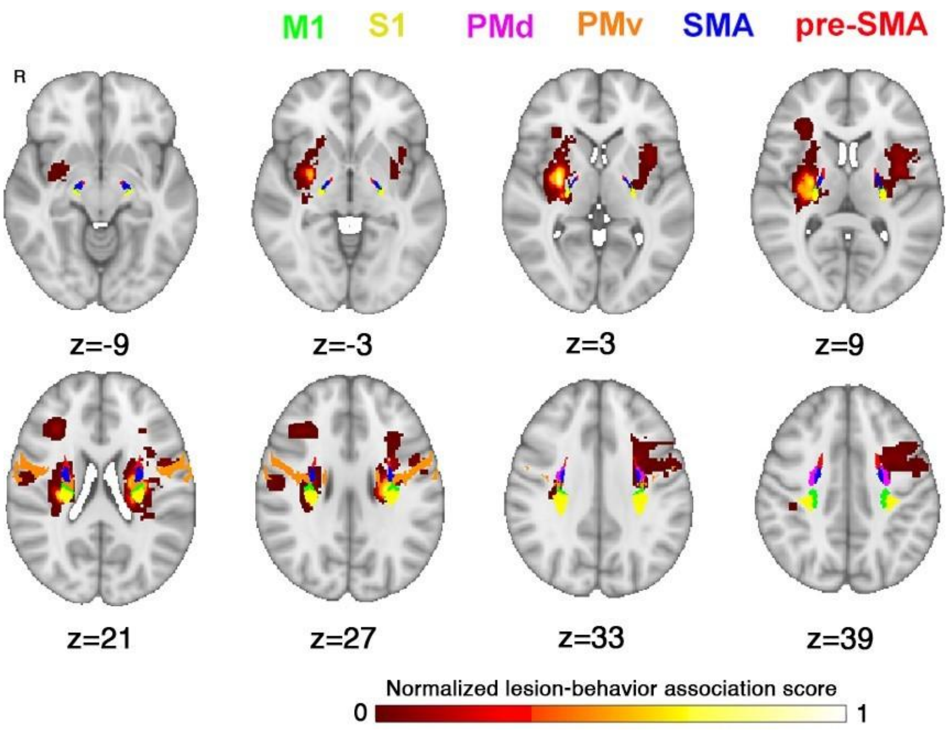


Supplementary Figure 10: Overlap between lesion-behaviour map and SMATT tracts. LBM map is plotted in hot colormap, SMATT tracts are plotted in their original colours (see Fig. 2).

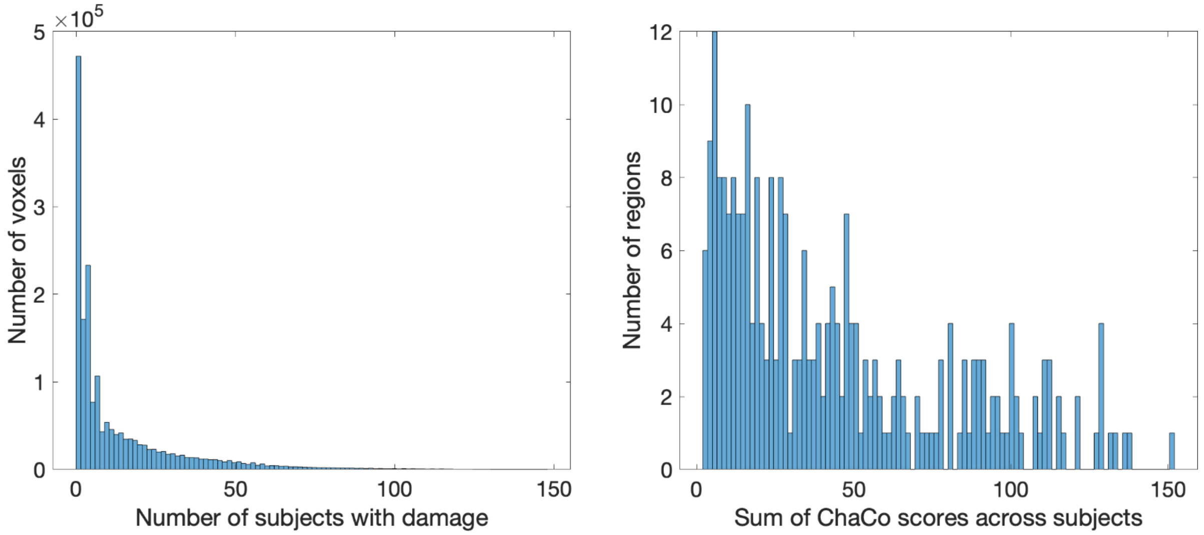


Supplementary Figure 11: Histogram of voxelwise lesion damage and regional (ChaCo scores) lesion damage. Left: Distribution of voxelwise lesion damage within a whole-brain mask. Most voxels are damaged in zero subjects. Right: Distribution of ChaCo scores across all subjects. Every region has some disconnection in at least one subject in the population.
